# DelSIEVE: cell phylogeny model of single nucleotide variants and deletions from single-cell DNA sequencing data

**DOI:** 10.1101/2023.09.09.556903

**Authors:** Senbai Kang, Nico Borgsmüller, Monica Valecha, Magda Markowska, Jack Kuipers, Niko Beerenwinkel, David Posada, Ewa Szczurek

## Abstract

With rapid advancements in single-cell DNA sequencing (scDNA-seq), various computational methods have been developed to study evolution and call variants on single-cell level. However, modeling deletions remains challenging because they affect total coverage in ways that are difficult to distinguish from technical artifacts. We present DelSIEVE, a statistical method that infers cell phylogeny and single-nucleotide variants, accounting for deletions, from scDNA-seq data. DelSIEVE distinguishes deletions from mutations and artifacts, detecting more evolutionary events than previous methods. Simulations show high performance, and application to cancer samples reveals varying amounts of deletions and double mutants in different tumors.

## Introduction

Cancer is a genetic disease driven by the accumulation of somatic mutations, resulting in highly heterogeneous cell populations [1–5]. The most common types of somatic mutations are single nucleotide variants (SNVs), followed by *deletions*, including point deletions, small deletions, and copy number aberrations. These events together can result in the activation of oncogenes and the inactivation of tumor suppressor genes, thus promoting tumor proliferation [2, 3, 5–8].

Single-cell DNA sequencing (scDNA-seq) technologies exhibit great potential for the analysis of intratumor genetic heterogeneity at the highest resolution of individual cells [9–12]. However, these technologies typically suffer from a low signal-to-noise ratio. Most rely on whole-genome amplification (WGA) before sequencing [12–16], with non-scWGA methods either not providing enough coverage to call SNVs [17] or only sequencing a panel of genes instead of whole genome or exome [18]. The WGA step introduces several biases in the sequencing data, including an uneven coverage of the genome, amplification errors, and allelic bias, where one of the paternal or maternal alleles is over- or underrepresented. Importantly, allelic bias can sometimes result in allelic (ADO) or locus dropout (LDO), where one or both alleles fail to be amplified [12–14].

Several different methods for calling SNVs from scDNA-seq data have been proposed. For instance, Monovar [19] employs consensus filtering based on the data from multiple cells, while other methods [20–22] leverage phase information from germline single nucleotide polymorphisms. The called SNVs are typically used for the reconstruction of the cell phylogeny [23–30]. As the cell phylogeny can be informative for SNV calling, SCIPhI [31] and our more recent model SIEVE [32] jointly infer the cell phylogeny and call SNVs.

However, the majority of methods that model the cell phylogeny or call SNVs from scDNAseq data do not account for deletions and consider only diploid genotypes during tumor evolution. The difficulty of accurate modeling of SNVs in the presence of deletions arises because the effects of deletions, back mutations, double mutants, allelic imbalance, and dropouts on sequencing data are often hard to distinguish. For example, several events might be the cause if only reads supporting the alternative nucleotide are observed. Assuming that one of the alleles has such alternative nucleotide, the other allele could either be deleted during evolution, be dropped out during amplification, or be mutated to exactly the same alternative nucleotide.

To address these ambiguities, methods such as SCARLET [33] or SCIPhIN [34] leveraged the idea that deletions occur along the cell phylogeny and thus groups of related cells should share the same deletions. However, these methods are unable to identify important evolutionary events such as double mutants (mutations affecting both alleles at a variant site) and do not fully exploit the information conveyed by sequencing coverage.

We reasoned that combining the information encoded in the raw read counts, especially the signal in sequencing coverage, and the phylogenetic relations among cells should result in more accurate inference of phylogenetic trees and variants in the presence of deletions. Indeed, despite the inherent noise in scDNA-seq data, it is expected that the sequencing coverage is proportional to the number of sequenced alleles and should provide information on the loss of alleles. On the other hand, the cell phylogeny should help to tell if the loss occurs during evolution or due to technical artifacts.

Here we introduce DelSIEVE (deletions enabled SIngle-cell EVolution Explorer), a statistical phylogenetic model that leverages both the signal from cell phylogeny and the coverage information, and explicitly accounts for the effect that deletions have on mutated sites. DelSIEVE can call seven different genotypes that not only include single or double mutants, but also single or double deletions, and is able to discern those from technical events such as ADO or LDO. Thanks to this increased expressive power, DelSIEVE is able to discern 28 types of genotype transitions, associated with 17 types of mutation events, many more than any existing method.

## Results

### Overview of the DelSIEVE model

DelSIEVE takes as input raw read counts for all four nucleotides for each cell *j ∈* {1*, …, J* } at each candidate site *i ∈* {1*, …, I*} in the form of the read counts of three alternative nucleotides with values in descending order, together with the total sequencing coverage (Figure 1a).

**Figure 1:**
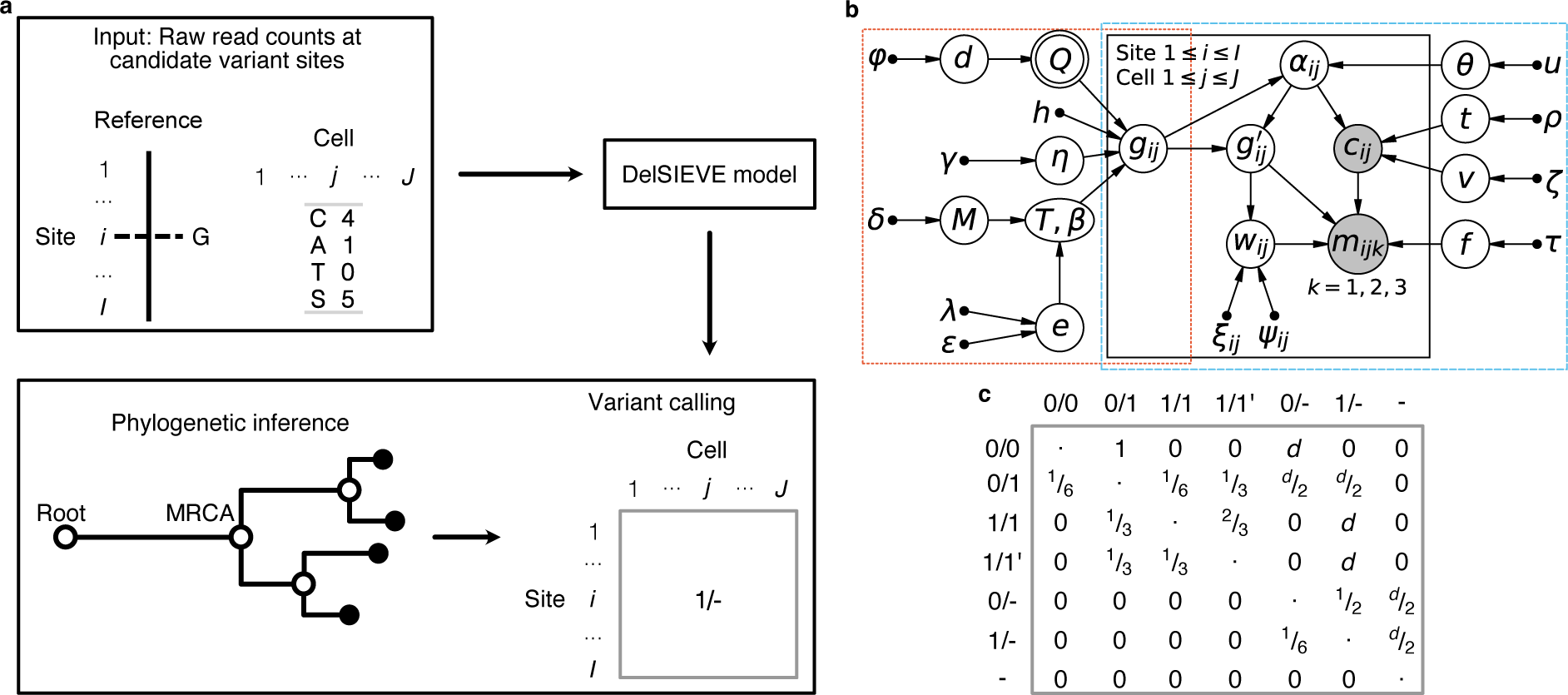
Overview of the DelSIEVE model. a. Analysis workflow of DelSIEVE with an example of input data. At candidate variate site *i ∈* {1*, …, I*}, the reference nucleotide is G. For cell *j ∈* {1*, …, J* } at site *i*, observed are the sequencing depth of 5 (marked by *S*) as well as read counts for nucleotide *C* being 4 and *A* being 1. DelSIEVE first infers from the input data the cell phylogeny, based on which the genotype state of each node in the tree is then determined through maximum likelihood estimation. For instance, 1/- is inferred as the genotype state of cell *j* at site *i*. **b** Probabilistic graphical model of DelSIEVE. The orange frame shows the part corresponding to the statistical phylogenetic model, and the blue frame encloses the part corresponding to the model of raw read counts. Shaded circular nodes represent observed variables, while unshaded circular nodes represent hidden random variables. Nodes with double circles are deterministic random variables, meaning that they are fixed once the values of their parent nodes are determined. Small black dots correspond to fixed hyper parameters. Arrows denote local conditional probability distributions of child nodes given parent nodes. **c** Instantaneous transition rate matrix of the statistical phylogenetic model. The hidden random variable *d* is the deletion rate, measured relatively to the mutation rate. The elements in the diagonal of the matrix are denoted by dots, and have negative values opposite to the sum of the other entries in the same row, ensuring that the sum of each row equals zero.

From that input data, the model first infers a tree phylogeny, which incorporates a trunk between the root (a normal cell) and the most recent common ancestor (MRCA) of the sampled tumor cells. DelSIEVE operates in a genotype state space that accounts both for SNVs and deletions of candidate variant sites. Specifically, apart from genotypes that were previously modeled by SIEVE: 0*/*0 (*wildtype*), 0*/*1 (*single mutants*), 1*/*1 (*double mutants*, where the two alternative nucleotides are the same) and 1*/*1^′^ (*double mutants*, where the two alternative nucleotides are different), DelSIEVE additionally considers 0/- (*reference-remaining single deletion*), 1/- (*alternative-remaining single deletion*) and - (*double deletions*). Here, 0, 1, 1^′^ and – represent the reference nucleotide, an alternative nucleotide, a second alternative nucleotide different from that denoted by 1, and deletions, respectively. The genotype state of each node in the tree is inferred using maximum likelihood estimation. As an effect, DelSIEVE is able to discern 28 types of genotype transitions, which we categorize into 17 different mutation events (eight more than SIEVE; see Section Mutation event classification and Table 3). These genotype transitions include 12 that were already previously modeled by SIEVE, and are complemented by 16 transition events associated to deletions.

The power of DelSIEVE lies in its probabilistic graphical model, where the hidden variable describing the genotype for site *i* in cell *j*, denoted *g_ij_*, is used as the bridge between the statistical phylogenetic model and the model of raw read counts (Figure 1b). The model accounts for the possible mutations using a deletion-aware instantaneous transition rate matrix (Meth-ods; Figure 1c). DelSIEVE employs a Dirichlet-Multinomial distribution to model the raw read counts for all nucleotides, and models the sequencing coverage using a negative binomial distribution, dependent on the number of alleles which can change due to deletions (see Methods for a detailed description).

### DelSIEVE accurately calls deletions

We first used simulated data to benchmark one of DelSIEVE’s advantages, namely calling deletions (see Section Simulation design in Additional file 1: Supplementary notes). To our knowledge, DelSIEVE is the only method that can differentiate alternative-remaining single deletion (genotype 1/-), reference-remaining single deletion (0/-), and double deletions (-), and thus it was not compared to any other method for these tasks.

For calling alternative- and reference-remaining single deletions, DelSIEVE achieved F1 scores with medians *≥* 0.87 and *≥* 0.76, respectively, when the data was of medium or high coverage quality (with high mean and low or medium variance of coverage; Figure 2a, b). For calling these two genotypes, the corresponding recall of DelSIEVE has medians *≥* 0.72 and *≥* 0.62 (Additional file 1: Figure S1a, c), the precision medians *≥* 0.96 and *≥* 0.97 (Additional file 1: Figure S1b, d), and the false positive rate (FPR) medians *≈* 0 (Additional file 1: Figure S2a, b). These results show that DelSIEVE can correctly and reliably identify most of the alternative- and reference-remaining single deletions.

**Figure 2.**
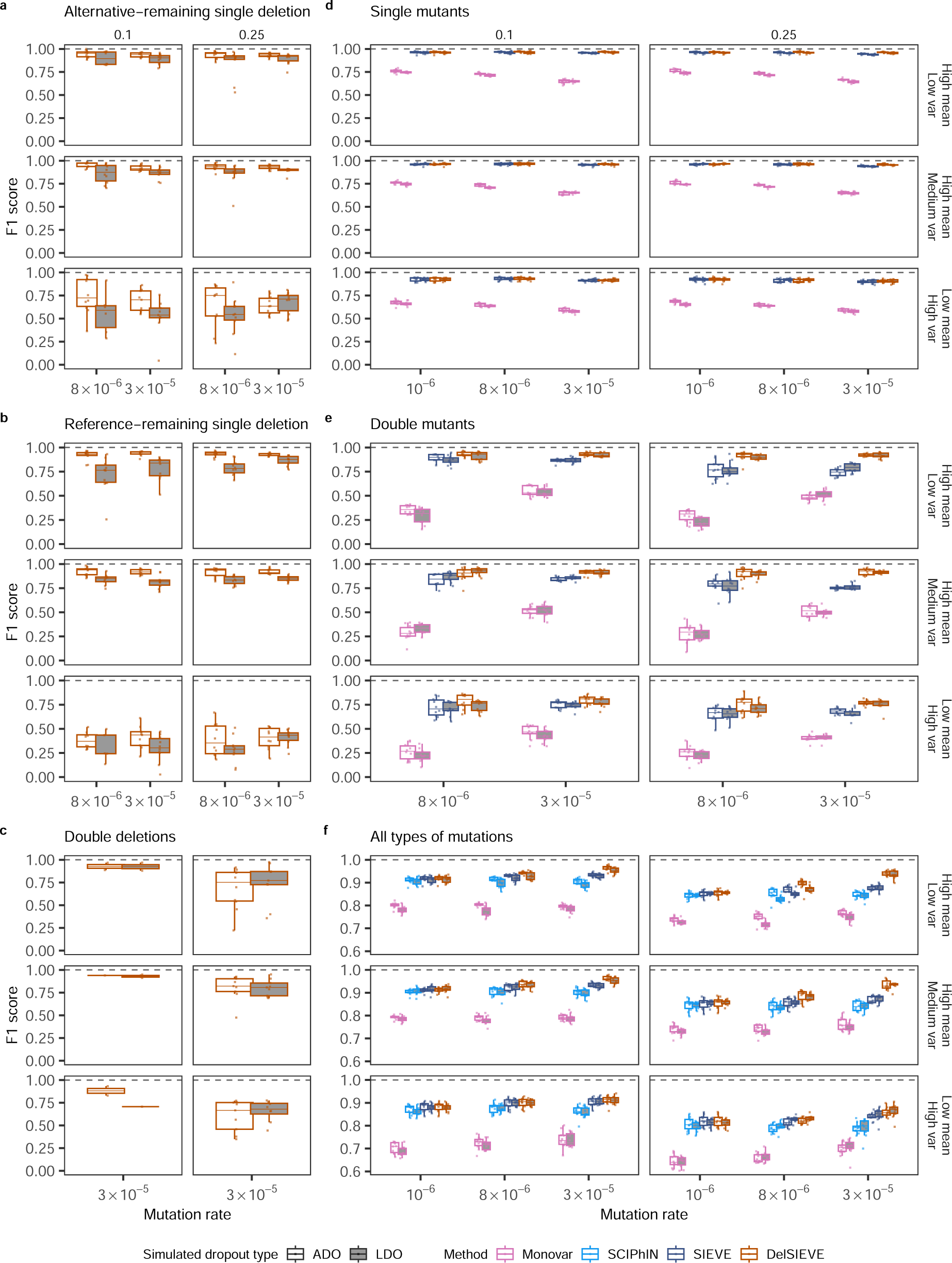
***(previous page)*: F1 score for the benchmark of the DelSIEVE model.** Varying are the mutation rate (the horizontal axis), the relative deletion rate (the vertical strip), the coverage quality (the horizontal strip), and the simulated dropout type (the shaded or blank boxes). Each simulation is repeated *n* = 10 times, with each repetition denoted by colored dots. The gray dashed lines represent the optimal values of each metric. Box plots comprise medians, boxes covering the interquartile range (IQR), and whiskers extending to 1.5 times the IQR below and above the box. Data points were removed if the proportion of simulated ground truth was less than 0.1%. Both DelSIEVE and SIEVE were configured to match the dropout mode (ADO or LDO) employed during the simulation process. **a-e**, Box plots of the F1 score for calling alternative-remaining single deletion (**a**), reference-remaining single deletion (**b**), double deletions (**c**), single mutants (**d**), and double mutants (**e**). The results in **c** when mutation rate was 8 *×* 10*^−^*^6^ were omitted as very few double deletions were generated (less than 0.2%; see Section Simulation design in Additional file 1: Supplementary notes). **f**, Box plots of the F1 score for calling all types of mutations considered in **a-e**.

When the data was of low coverage quality (low mean and high variance of coverage), the F1 score medians of DelSIEVE dropped to *≥* 0.55 and *≥* 0.29 for calling alternative- and referenceremaining single deletions, respectively (Figure 2a, b). The low quality of the data affected more the calling of the reference-remaining single deletion (Additional file 1: Figure S1a-d), which was expected as such low coverage provided little information for this task. Furthermore, the FPR of DelSIEVE was still *≈* 0 for the low quality data.

When calling double deletions, DelSIEVE obtained high F1 scores medians *≥* 0.75 (Figure 2c). Its performance decreased as the deletion rate increased or the coverage quality of the data decreased (Figure 2c, Additional file 1: Figure S1e, f), but the FPR was kept at a negligible level (*≈* 0; see Additional file 1: Figure S2c).

We observed that the performance of DelSIEVE only slightly decreased in the presence of LDO, in comparison to the results obtained when it was applied to data simulated under the ADO mode. Given that DelSIEVE explicitly models the sequencing coverage, it was anticipated that data simulated under the LDO mode would introduce additional uncertainties to the inference.

### DelSIEVE reliably identifies mutations in the presence of deletions

We next assessed DelSIEVE’s performance in calling single and double mutants against Monovar and SIEVE (Figure 2d, e, Additional file 1: Figure S3, S4).

Regarding calling single mutants, DelSIEVE and SIEVE performed comparatively well (minimum median F1 score 0.9), and outperformed Monovar (minimum median F1 score 0.58; see Figure 2d). This advantage can be due to the fact that in contrast to Monovar, DelSIEVE and SIEVE consider the cell phylogeny during variant calling. As the mutation rate increased, the recall of both DelSIEVE and SIEVE slightly increased (Additional file 1: Figure S3a), while the precision slightly decreased (Additional file 1: Figure S3b), resulting in relatively constant F1 scores. In contrast, Monovar experienced a decrease in both recall and precision as the mutation rate increased (Additional file 1: Figure S3a, b). Moreover, DelSIEVE and SIEVE had comparable recall (Additional file 1: Figure S3a), while DelSIEVE showed higher precision (Additional file 1: Figure S3b) and lower FPR (Additional file 1: Figure S4a) than SIEVE, especially when the mutation rate was high (3 *×* 10*^−^*^5^). We speculate that this might be because SIEVE has to interpret the evident signal of deletions as ADO or LDO events occurring in addition to mutations.

Additionally, as the mutation rate increased, the FPR of all methods also increased (Additional file 1: Figure S4a). It was noteworthy that, when the mutation rate was high (*≥* 3*×*10*^−^*^5^), DelSIEVE and SIEVE had slightly higher FPR than Monovar for calling single mutants (Addi-tional file 1: Figure S4a). However, this loss was negligible compared to SIEVE and DelSIEVE’s advantage over Monovar when considering precision, recall, and F1 score.

In the task of calling double mutants, Monovar obtained minimum median F1 scores of 0.21, while SIEVE and DelSIEVE exhibited much better performance with minimum median F1 scores 0.65 and 0.93, respectively (Figure 2e). More specifically, DelSIEVE and SIEVE had a comparable recall (Additional file 1: Figure S3c), but the former reached higher precision (minimum medians 0.75 and 0.61, respectively; see Additional file 1: Figure S3d). Again, this discrepancy in performance could be due to SIEVE’s inclination to explaining deletions as dropout events occurring on top of double mutants.

DelSIEVE also had the lowest FPR (*≈* 0) (Additional file 1: Figure S4b). These findings highlighted the superior accuracy of DelSIEVE in identifying double mutants in the presence of deletions. On top of that, the slight advantage of Monovar over methods incorporating phylogeny for calling single mutants was not observed for calling double mutants. In contrast, Monovar had a significantly elevated FPR in this task compared to all other methods.

### DelSIEVE outperforms alternative models in variant calling, regardless of the variant type

To compare to one more predecessor model, SCIPhIN, which does not distinguish among single and double mutants, as well as alternative-remaining single deletion, reference-remaining single deletion and double deletions, we considered all genotypes other than wildtype as general “mutations” and computed the related performance metrics (see Sections Variant calling and phylogenetic accuracy in Additional file 1: Supplementary notes).

Overall, Monovar was outperformed by the other three methods (Figure 2f, Additional file 1: Figure S5a-c), which had similar performance when the mutation rate was low (10*^−^*^6^). As the mutation rate increased, DelSIEVE performed better than SIEVE and SCIPhIN (Figure 2f). Specifically, DelSIEVE had higher recall compared to SIEVE and SCIPhIN (Additional file 1: Figure S5a), with similar precision and FPR (Additional file 1: Figure S5b-c). With the increase of the relative deletion rate and the decrease of the coverage quality, the performance of all methods slightly dropped. The dropout mode under which the data was simulated seemed to have an insignificant effect on all methods, except for the precision and FPR of Monovar, which were worse under the LDO mode (Additional file 1: Figure S5b, c).

### DelSIEVE can identify ADO and LDO

We then evaluated DelSIEVE’s performance in calling ADO and LDO against SIEVE (Figure 3, Additional file 1: Figure S6, S7), which are the only two methods that can infer these events. Though unsupported originally, we implemented the LDO mode in SIEVE for this comparison (see Section Configurations of methods in Additional file 1: Supplementary notes).

**Figure 3:**
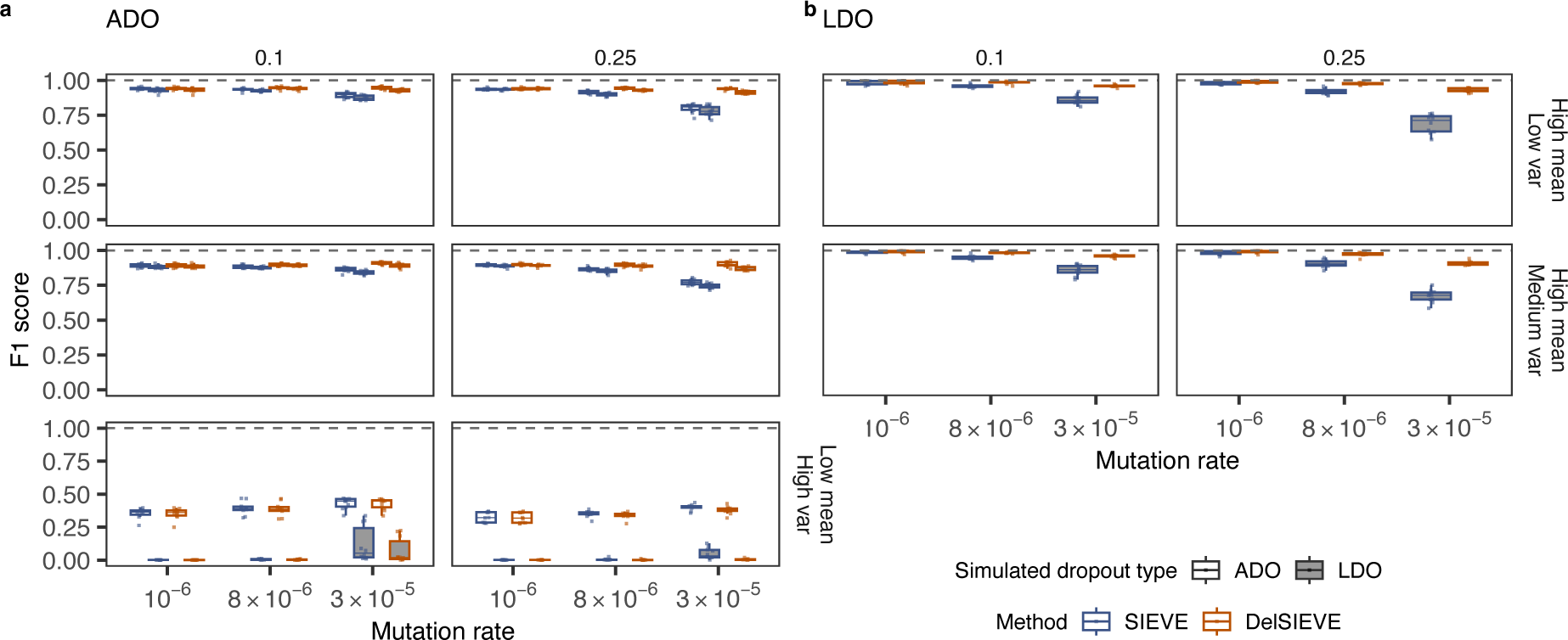
F1 score for the benchmark of calling ADO and LDO. Varying are the mutation rate (the horizontal axis), the relative deletion rate (the vertical strip), the coverage quality (the horizontal strip), and the simulated ADO type (the shaded or blank boxes). Each simulation is repeated *n* = 10 times, with each repetition denoted by colored dots. The gray dashed lines represent the optimal values of each metric. Box plots comprise medians, boxes covering the interquartile range (IQR), and whiskers extending to 1.5 times the IQR below and above the box. Both DelSIEVE and SIEVE were configured to match the dropout mode (ADO or LDO) employed during the simulation process. **a-b**, Box plots of the F1 score for calling ADO (**a**) and LDO (**b**). The F1 scores were unavailable in **b** when data was of low coverage quality due to unavailable precision. The results of calling LDO for data simulated with ADO are not available in (**b**), as both models were configured for the same dropout mode as used in the simulated data and were not able to call LDO in this case.

ADO calling was affected by the coverage quality. When the data was of medium or high coverage quality, DelSIEVE reached a minimum median F1 score of 0.9, higher than SIEVE (0.77; see Figure 3a). The performance of DelSIEVE remained consistent regardless of changes in the mutation and deletion rates, in contrast to SIEVE. This was anticipated because higher mutation or deletion rates resulted in an increased number of deletions being generated. DelSIEVE was capable of differentiating deletions from ADOs, while SIEVE wrongly accounted for deletions as ADOs occurring on top of single or double mutants. This behavior reduced SIEVE’s recall and precision, and increased FPR (Additional file 1: Figure S6a, b, Additional file 1: Figure S7a), as when calling single and double mutants (see Section DelSIEVE reliably identifies mutations in the presence of deletions). The performance of DelSIEVE and SIEVE in calling ADO declined when the data had low coverage quality (Figure 3a, Additional file 1: Figure S6a, b, Additional file 1: Figure S7a). This decrease in performance was further exacerbated when the data was simulated under the LDO mode.

When calling LDOs from data of medium or high coverage quality, DelSIEVE showed a minimum median F1 score of 0.91, higher than SIEVE did (0.68; see Figure 3b). Specifically, DelSIEVE and SIEVE were comparable in terms of recall (Additional file 1: Figure S6c), but DelSIEVE had a higher precision and lower FPR as the mutation and deletion rates increased (Additional file 1: Figure S6d, Additional file 1: Figure S7b). However, when the data was of low coverage quality, both methods reported no LDO, resulting in zero recall and FPR as well as not available precision and F1 score values.

### DelSIEVE estimates accurate cell phylogenies

We further benchmarked DelSIEVE’s performance in reconstructing the cell phylogeny against SiFit, SCIPhIN and SIEVE (Additional file 1: Figure S8). To measure phylogenetic accuracy, we used the branch score (BS) distance, which considers both tree topology and branch lengths and the normalized Robinson-Foulds (RF) distance, which only considers the tree topology (see Section Variant calling and phylogenetic accuracy in Additional file 1: Supplementary notes). The results of SCIPhIN were excluded in the computation of the BS score as it does not estimate branch lengths.

DelSIEVE and SIEVE outperformed SiFit when branch lengths were considered, showing the advantage of correcting the acquisition bias (Additional file 1: Figure S8a). Moreover, all methods tended to overestimate branch lengths when the mutation rate was higher (*≥* 8 *×* 10*^−^*^6^). The performance of DelSIEVE and SIEVE in topology reconstruction was similar (maximum median normalized RF distance 0.29 and 0.28, respectively), and better compared to SiFit (maximum median normalized RF distance 0.37) and SCIPhIN (0.33; see Additional file 1: Figure S8b), especially when the mutation rate increased. DelSIEVE and SIEVE were robust to variations in mutation rates in comparison to SiFit and SCIPhIN, while the performance of all methods declined as the coverage quality decreased. The high performance of DelSIEVE in variant calling and phylogenetic reconstruction is likely due to the benefit of sharing information between these two tasks.

### The dropout mode configuration of DelSIEVE has negligible effect on performance

The previous results were obtained with DelSIEVE configured to match the dropout mode (ADO or LDO) employed during the simulation process. To investigate the effects of model misspecification, we further ran DelSIEVE (and, for completeness, where possible, also SIEVE) under a dropout mode different from that used to simulate the data (see Section Configurations of methods in Additional file 1: Supplementary notes).

The configuration of the dropout mode, regardless of that used in the simulated data, did not significantly affect DelSIEVE’s calling of deletions (Additional file 1: Figure S9a-c), or DelSIEVE’s and SIEVE’s calling of single and double mutants (Additional file 1: Figure S9d, e). We also observed that for simulated data of high coverage quality under the ADO mode, the dropout mode of DelSIEVE and SIEVE did not affect ADO calling (Additional file 1: Figure S10). However, for data of the high coverage quality but under the LDO mode, it was favored to run those methods under the same dropout mode. On the contrary, when the data was of low coverage quality, it was favorable to run both methods under the ADO mode, regardless of that used to generate the data. Finally, the dropout configuration did not affect the phylogeny reconstruction of SIEVE and DelSIEVE (Additional file 1: Figure S11a-b), except for the high mutation rate and coverage quality for the BS score of DelSIEVE, where running under ADO mode slightly increased the estimated branch lengths (Additional file 1: Figure S11a). Since the real dataset resembles the low coverage quality data, running DelSIEVE under ADO mode should have a negligible effect on the tree reconstruction.

Given that the LDO versus ADO mode configuration affects the model’s performance only slightly and given that LDOs are relatively rare compared to ADOs, we ran DelSIEVE in ADO mode for the analysis of the real datasets discussed below.

### DelSIEVE identifies deletions in triple negative breast cancer (TNBC) cells

We applied DelSIEVE to real scDNA-seq datasets previously analyzed using SIEVE [32] (see Section Configurations of methods in Additional file 1: Supplementary notes). For the single-cell whole-exome sequencing (scWES) dataset TNBC16, containing data for 16 cells [35], DelSIEVE reported a maximum clade credibility cell phylogeny with a long trunk, and with high posterior probabilities for most nodes (Figure 4, Additional file 1: Figure S12). The cell phylogeny was very similar to that reported by SIEVE, with the normalized RF and BS distances being 0.07 and 3.88 *×* 10*^−^*^6^, respectively.

**Figure 4:**
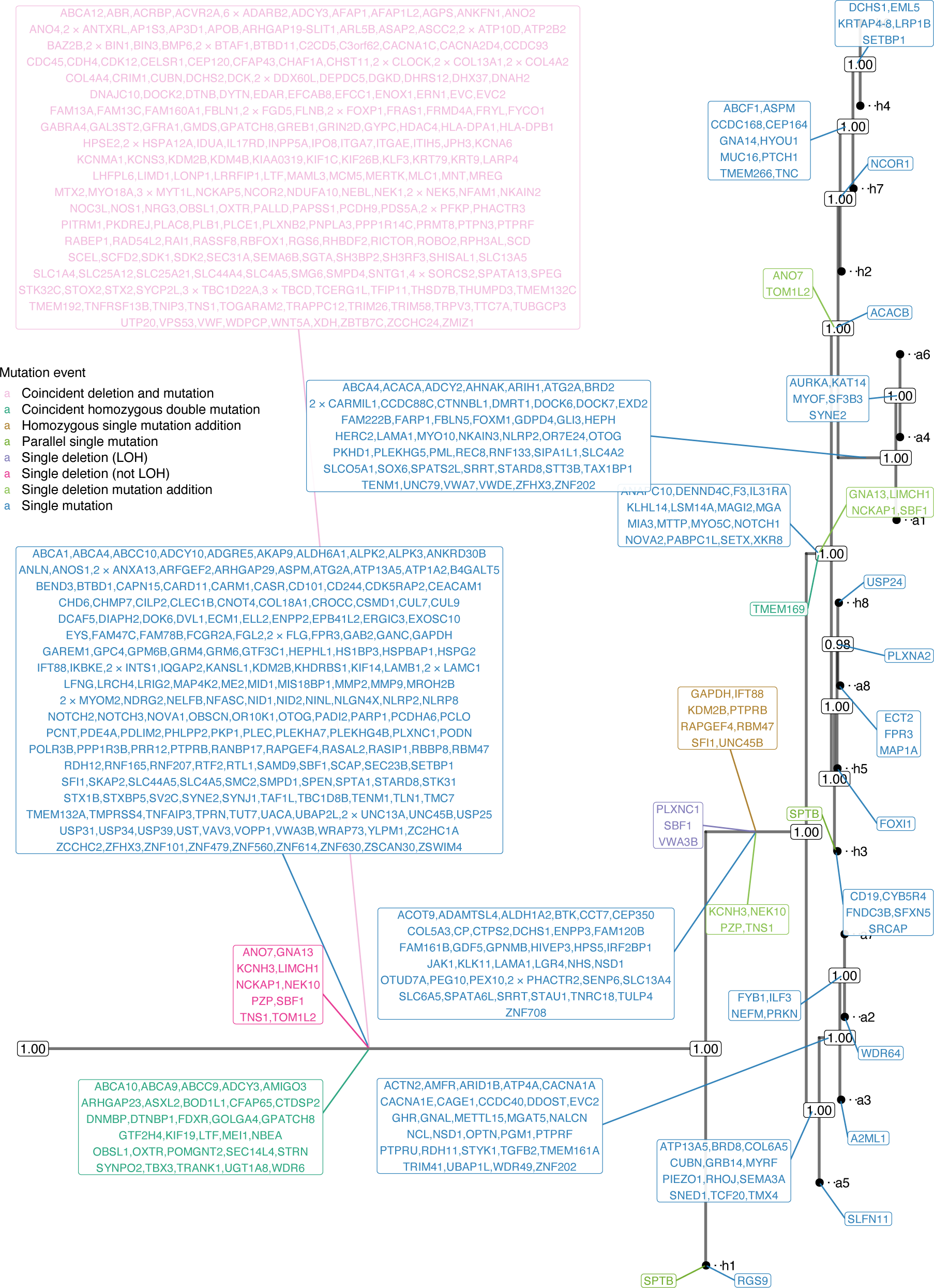
Results of phylogenetic inference for the TNBC16 dataset. Shown is DelSIEVE’s maximum clade credibility tree. Tumor cell names are annotated to the leaves of the tree. The numbers at each node represent the posterior probabilities (threshold *p >* 0.5). At each branch, depicted in different colors are non-synonymous genes that are either TNBC-related single mutations (in blue) or other mutation events (in other colors; see the legend).

DelSIEVE identified the same types of mutation events as reported by SIEVE, except for back mutations. In terms of numbers, DelSIEVE explained the same data with fewer point mutations. Specifically, DelSIEVE identified 31 coincident homozygous double mutations (transitions from 0/0 to 1/1; 44 for SIEVE), eight homozygous single mutation additions (from 0/1 to 1/1; nine for SIEVE), and two parallel single mutations (from 0/0 to 0/1 that occurred more than once in the tree; same for SIEVE). SIEVE found seven single back mutations (from 0/1 to 0/0; *BRD8*, *COL6A5*, *GRB14*, *MYRF*, *RHOJ*, *SEMA3A*, *TMX4*), which were gained in the trunk and lost afterward in the branch leading to the a2/a3/a5/a7 clade. In contrast, DelSIEVE identified those as unique mutations that occurred only in the ancestor of this clade.

In addition, DelSIEVE identified several deletions, including a large number of 245 coincident deletions and mutations (from 0/0 to 1/-), three single deletions which could be categorized as LOH (from 0/1 to 0/- or 1/-, or from 1*/*1^′^ to 1/-), ten single deletions which were not LOH (from 0/0 to 0/-, or from 1/1 to 1/-), and finally ten single deletion mutation additions (from 0/- to 1/-). For instance, DelSIEVE inferred that gene *NEK1* and *NEK5*, which had been reported to be related to breast tumors [36], experienced both a deletion and a mutation on the trunk, resulting in all sequenced cells having genotype 1/-. Another gene, *LIMCH1*, known to be related to TNBC [37], had an allele deleted first on the trunk (genotype changed from 0/0 to 0/-), and then the remaining allele mutated for a subgroup of cells (genotype changed from 0/- to 1/-). The substantial amount of evolutionary events related to deletions highlights the importance of the extended functionality of DelSIEVE as compared to SIEVE.

In total, DelSIEVE identified 5,893 variant sites, close to the 5,895 variant sites reported by SIEVE (Figure 5). Among the 683 sites inferred by DelSIEVE that contain deletions (mostly 1/-; 11.6% of all variant sites), 377 were previously determined according to SIEVE to have double mutants and the remaining 306 to have a single mutant genotype. This observation was in accordance with the simulation results, where SIEVE tended to explain deletions as dropout events within single and double mutants. The proportions of different genotypes called by DelSIEVE and SIEVE are summarized in Additional file 1: Table S4 (same for the following datasets).

**Figure 5:**
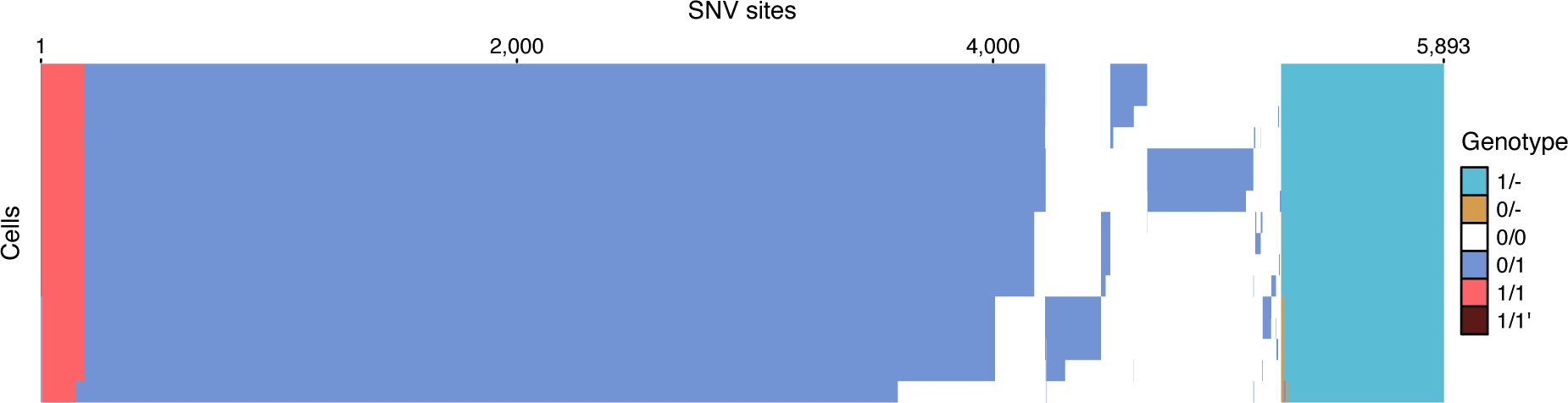
Results of variant calling for the TNBC16 dataset. Cells in the rows are in the same order as that of leaves in the phylogenetic tree in Figure 4.

### DelSIEVE identifies rare mutations in colorectal cancer (CRC) cells

We then applied DelSIEVE to a scWGS dataset, CRC28 [32], containing data for 28 single cells coming from three biopsies: tumor proximal (TP; with nine cells), tumor distal (TD; with seven cells) and tumor central (TC; with 12 cells). The estimated cell phylogeny was supported by high posterior probabilities with a long trunk (Additional file 1: Figure S13, S14), which was similar to that reported by Kang *et al.* using SIEVE (the normalized RF and the BS distances were 0.08 and 8.03 *×* 10*^−^*^7^, respectively). In particular the TP and TD subclones also formed sister clades in the tree reported by DelSIEVE, with the TC subclone forming a separate clade, suggesting regular tumor growth and limited cell migration.

Similarly to SIEVE, DelSIEVE identified mutations in known CRC driver genes, for instance, *APC*, and of genes related to the metastatic progression of CRC, such as *ASAP1* and *RGL2* on the trunk of the tree. However, DelSIEVE identified more mutation events than SIEVE, including two coincident deletions and mutations, one single deletion which was not LOH, and one single deletion mutation addition. For example, DelSIEVE identified that *ACSL5*, potentially related to intestinal carcinogenesis [38], underwent a somatic deletion of one allele (genotype changed from 0/0 to 0/-) on the trunk and a mutation to the remaining allele (genotype changed from 0/- to 1/-) for the most recent common ancestor of TP and TD subclones.

DelSIEVE identified the same number of variant sites as SIEVE (8,029; see Additional file 1: Figure S15), in which 13 sites contained deletions (mostly 1/-; 0.16% of all variant sites). According to SIEVE, nine of those sites were inferred to have double mutants and four to have single mutants. Overall, both DelSIEVE and SIEVE found very few mutation events that were not single mutations. The contrasting results obtained by DelSIEVE for TNBC16 (with multiple deletions) compared to CRC28 (only few deletions), underscored an important feature of the method. While DelSIEVE employs a sophisticated and expressive modeling approach, it primarily relies on the data for the inference, without “overcalling” deletions.

### DelSIEVE identified rare somatic mutations in CRC samples mixed with normal cells

We finally analyzed another scWES dataset, CRC48 (CRC0827 in [39]), with cells collected at three anatomical locations: adenomatous polyps (containing 13 normal cells), cancer tissue 1 (17 cells) and cancer tissue 2 (18 cells). DelSIEVE pinpointed two tumor subclones, associated with their anatomical locations, each subclone containing exactly the same cells as found by Kang *et al.* using SIEVE (Additional file 1: Figure S16, S17). The rest of the cells collected from tumor biopsies were clustered together with cells from adenomatous polyps, suggesting that they might be normal cells residing inside cancer tissues, as pointed out by both the original study [39] and Kang *et al.* [32]. There were some distinctions between the cell phylogenies reported by DelSIEVE and SIEVE, with normalized RF and BS distances being 0.33 and 1.99 *×* 10*^−^*^6^, respectively. This discrepancy is higher than observed for the previous datasets, and might be due to the overall lower signal level in the data. Indeed, the CRC48 dataset has a substantially lower ratio between the number of candidate variant sites and the number of cells (707*/*48 *≈* 14.7) compared to TNBC16 (5912*/*16 = 369.5) and CRC28 (8470*/*28 = 302.5), and therefore contains potentially less phylogenetic information.

DelSIEVE identified many single mutations on the branch leading to the two tumor subclones, including a reported CRC driver mutation in gene *SYNE1* [40], as well as a mutation related to DNA mismatch repair, in gene *MLH3* [41]. DelSIEVE also found two parallel single mutations (*CHD3* and *PLD2*). Furthermore, DelSIEVE identified only one site containing deletions (among 679 variant sites, and only 0/-; see Additional file 1: Figure S18), which was previously inferred by SIEVE to have a single mutant genotype.

### Sequencing coverage agrees with deletion sites identified by DelSIEVE

To further validate the ability of DelSIEVE to reliably call deletions, we inspected whether the sites identified as deleted in the analyzed datasets had lower coverage than sites lacking deletions. We next compared the strength of the coverage reduction effect on deleted sites with the results from dedicated copy number calling methods, Sequenza [42] (applicable to bulk sequenced samples from TNBC16, TD subclone from CRC28, and cancer tissue 1 and 2 from CRC48), as well as Ginkgo [43] (Figure 6) (applicable to WGS of single cells in CRC28). The comparison was performed only for the candidate variant sites, and the raw sequencing coverage was scaled by the corresponding size factors of the single cells.

**Figure 6.**
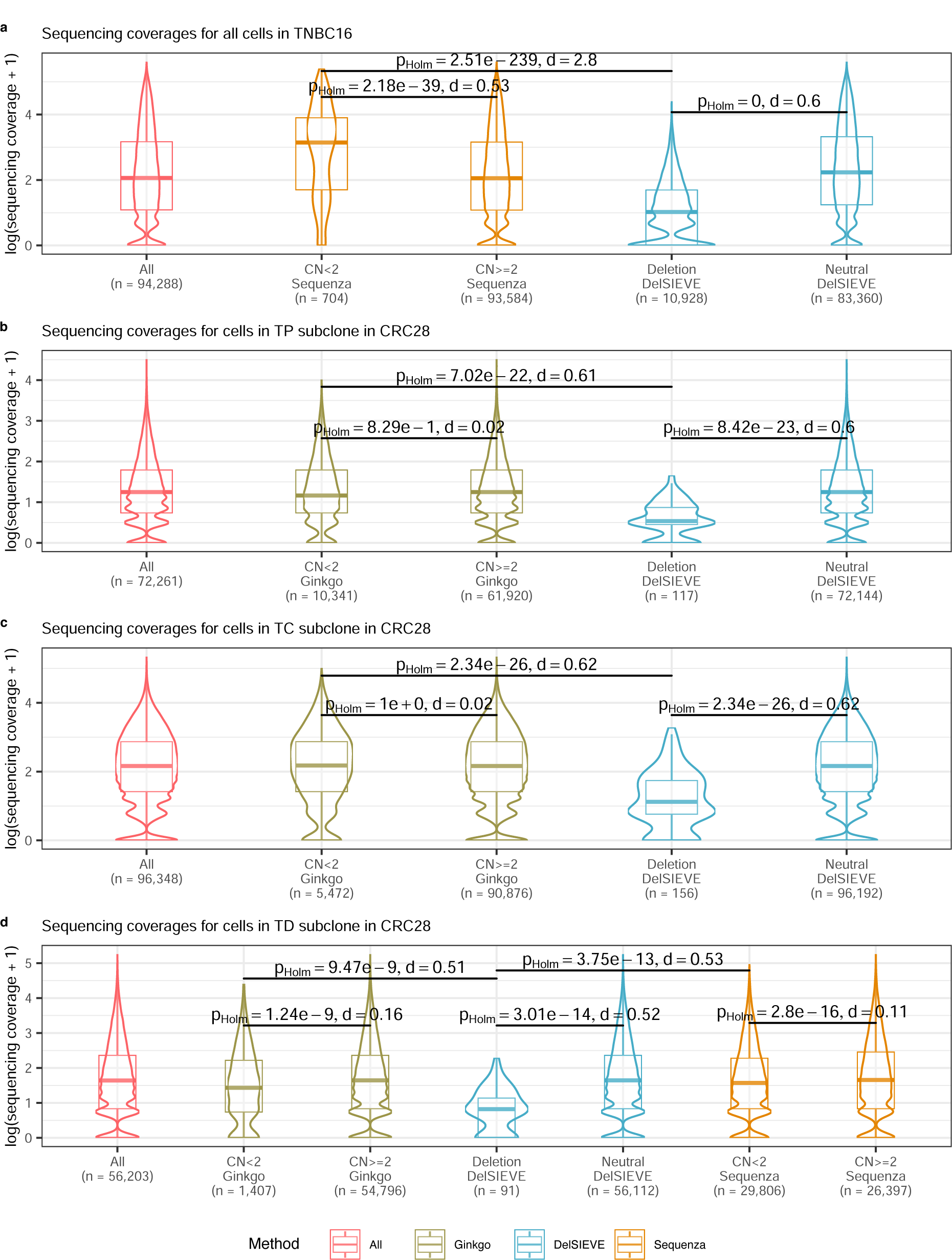
***(previous page)*: Comparison of sequencing coverage on clone (for TNBC16) and subclone (for CRC28) level.** Compared were the sites shared in the output of DelSIEVE, Sequenza [42] and Ginkgo [43], if available. For Sequenza and Ginkgo, sites were divided into two groups with copy number (CN) *<* 2 and *≥* 2. For DelSIEVE, sites were also divided into two groups, one with deletions, the other copy neutral. Sequencing coverage transformed with log *p*1 across all cells in the clone or subclone at all sites were plotted for reference. In each group, the violin and the box plots matched the color of the method. The total number of data points in each group was marked with *n* on the horizontal axis. Box plots comprise medians, boxes covering the interquartile range (IQR), and whiskers extending to 1.5 times the IQR below and above the box. Within- and between-group comparisons were conducted between CN *<* 2 and *≥* 2 of Sequenza and Ginkgo, between deletions and copy neutral of DelSIEVE, and between deletions of DelSIEVE and CN *<* 2 of Sequenza and Ginkgo. Each comparison was conducted on the sequencing coverage on the original scale, showing the result of two-sided Mann-Whitney U test, with the p-value corrected by Holm–Bonferroni method, and the absolute value of the effect size (Cohen’s d). **a-d**, Comparison of sequencing coverage for all cells in TNBC16 (**a**) as well as in TP (**b**), TC (**c**), and TD (**d**) subclones in CRC28.

Since Sequenza was designed for bulk-seq data, we adjusted the resolution of DelSIEVE’s and Ginkgo’s results with Sequenza for the sake of comparison. To this end, a site in a given sample was called a variant site (a deletion for DelSIEVE, deletion or amplification for Gingko) if the method identified that variant in at least one cell from the sample. All other sites were considered neutral, where no cells had deletions or amplifications.

For the TNBC16 dataset the mean value of sequencing coverage in the group of sites with deletions identified by DelSIEVE (3.59) was significantly lower compared to the mean for sites without deletions (22.04), with effect size Cohen’s d = 0.6 (Figure 6a). In contrast, the mean coverage for 44 sites identified as containing deletions by Sequenza was 36.13, significantly larger than 19.78, the mean coverage for sites with amplifications (Cohen’s d = 0.53), controverting Sequenza’s copy number calls for the TNBC16 dataset. Furthermore, a direct comparison revealed that sites identified as deleted by DelSIEVE showed much lower coverage levels than those identified as deleted by Sequenza (Cohen’s d = 2.8).

For the CRC28 dataset and the TP subclone (Figure 6b) as well as the TC subclone (Figure 6c), the mean coverage at sites with deletions called by DelSIEVE was significantly lower than that from sites without deletions (Cohen’s d = 0.6 and 0.62 for TP and TC subclones, respectively). However, such differences were not significant for Ginkgo (Cohen’s d = 0.02 for both TP and TC subclones).

For the TD subclone (Figure 6d), DelSIEVE, Ginkgo and Sequenza had a lower mean coverage for sites with deletions compared to those without, where DelSIEVE exhibited a more evident distinction (Cohen’s d = 0.52) than Ginkgo and Sequenza (Cohen’s d = 0.16 and 0.11, respectively). Moreover, for sites containing deletions called by the three methods, DelSIEVE had the lowest mean coverage, which was significantly different from the mean coverages of Ginkgo and Sequenza (Cohen’s d = 0.51 and 0.53, respectively).

We also visualized across the entire genome the reported CNs of Ginkgo in all cells and of Sequenza in TD subclone (Additional file 1: Figure S19a-b). Based on the CNs called by Ginkgo, it was evident that the phylogenetic distance between TP and TD subclones was shorter than that between either of them and TC subclone, as in the tree reported by DelSIEVE (Additional file 1: Figure S13, S14). Moreover, although Sequenza inferred a majority of deletions, Ginkgo only inferred a small number of deletions, in accordance with the results of DelSIEVE, where only few deletions were identified.

Finally, for the CRC48 dataset, sites with and without deletions identified by DelSIEVE showed a pronounced mean coverage difference, for both cancer tissue 1 (Additional file 1: Figure S20; Cohen’s d = 0.39) and cancer tissue 2 (Additional file 1: Figure S21; d = 0.48). The mean coverage difference between sites identified as deleted or not by Sequenza was negligible for both subclones (Cohen’s d = 0.04 for cancer tissue 1; d = 0.09 for cancer tissue 2). Moreover, the mean coverage was much lower for sites identified to carry deletions by DelSIEVE than for sites identified as carrying deletions by Sequenza (Cohen’s d = 0.45 for cancer tissue 1; d = 0.51 for cancer tissue 2).

## Discussion

We present DelSIEVE, a statistical method designed to jointly infer deletions, SNVs, and the cell phylogeny from scDNA-seq data. Built upon SIEVE, which combines the inference of SNVs and the cell phylogeny, DelSIEVE takes a step forward by also considering point deletions. In a nutshell, DelSIEVE features a statistical phylogenetic model with genotypes relating both to deletions and to single and double mutants, a model of raw read counts allowing for both ADO and LDO, and a mechanism for acquisition bias correction for the branch lengths.

Deletions often play an essential role in tumor evolution. We have shown that SIEVE tends to explain deletions as a result of dropouts, and overestimates the amount of single and double mutants. Compared to SIEVE, DelSIEVE exhibits improved performance in terms of calling double mutants, while performing similarly in estimating the cell phylogeny and calling single mutants.

The difficulty of identifying deletions from scDNA-seq data is mainly due to the fact that dropouts and uneven coverage, prevalent in this type of data, can also decrease the observed coverage at a site. DelSIEVE is the only method capable of discerning detailed types of deletions such as alternative-remaining, reference-remaining and double deletions. DelSIEVE outperforms Monovar, SCIPhIN and SIEVE in variant calling on simulated data. With high enough coverage quality, DelSIEVE outperforms the only other approach, SIEVE, for ADO and LDO calling. When applied to three real scDNA-seq datasets from TNBC and CRC samples, which were previously analyzed using SIEVE, DelSIEVE identified rare deletions and double mutants in the CRC samples, akin to the results of SIEVE. However, for the TNBC dataset, DelSIEVE identified multiple deletions while revealing fewer single and double mutants compared to SIEVE, consistent with the benchmarking results.

A potential improvement to DelSIEVE would be to add the identification of insertions. Moreover, the current procedure for preselecting the candidate variant sites is limited to those sites that potentially contain nucleotide substitutions. To address this limitation, a possible enhancement would be to enable this procedure to preselect sites of tumor suppressor genes that are solely associated with deletions. The inclusion of these sites, which are known to elevate the risk of tumor development [3, 44], could further refine DelSIEVE’s utility in understanding tumorigenesis and potential therapeutic targets. Furthermore, an important assumption underlying DelSIEVE is that the genomic sites are independent for computational reasons. However, this assumption is violated for copy number aberrations (CNAs). Thus, DelSIEVE is designed only for the identification of point mutations, not for the detection of copy number changes in consecutive genomic regions.

Despite these limitations, DelSIEVE is one of the most sophisticated statistical phylogenetics models available. The expanded capabilities of DelSIEVE make it a valuable tool for unraveling complex genomic dynamics and understanding evolutionary relationships among cells.

## Conclusions

DelSIEVE is a novel probabilistic model that from raw read counts of scDNA-seq data jointly infers cell phylogeny and somatic variants, including SNVs and their deletions. We prove in our simulations that DelSIEVE is able to reliably differentiate several types of deletions and SNVs, while also reporting highly credible cell phylogenies. DelSIEVE can also call different types of dropout events, namely ADOs and LDOs, provided that the data is of enough quality, which is highly promising as the technology continues to advance. The application of DelSIEVE is not limited to tumors; the model can also be employed to investigate evolutionary dynamics in other tissue types.

## Methods

### Statistical phylogenetic model behind DelSIEVE

For the genotype state space *G* = {0*/*0, 0*/*1, 1*/*1, 1*/*1^′^, 0/-, 1/-, -} given for the DelSIEVE model, we define the instantaneous transition rate matrix *Q* as visualized in Figure 1c. We set the somatic mutation rate to 1 [45], where the relative measurements for the back mutation rate and deletion rate are ^1^*/*_3_ and *d*, respectively. Thus, *Q* is deterministic and depends on the value of the relative deletion rate *d*, namely *P* (*Q | d*) = 1. Each entry in *Q* represents the transition rate from the genotype in the row to that in the column during an infinitesimal time Δ*t*, while each row in *Q* sums up to 0. The continuous-time homogeneous Markov chain underlying *Q* is time non-reversible and reducible. For instance, genotypes that have both alleles present can transition to genotypes with one or both alleles lost, but not vice versa. To be specific, genotypes {0*/*0, 0*/*1, 1*/*1, 1*/*1^′^} and genotypes {0/-, 1/-} form two ergodic, transient communicating classes, while genotype {-} forms a closed communicating class. As a result, the limiting distribution of the Markov chain exists, where the value corresponding to genotype - is 1, while the others are 0.

Denote by *g_ij_* the hidden variable describing the genotype for site *i ∈* {1*, …, I*} in cell *j ∈* {1*, …, J* }. Based on the well-established theory of statistical phylogenetic models [45], the joint conditional probability of the genotype states of all sequenced cells at site *i*, namely ***g***^(^*^L^*^)^_*i*_, is

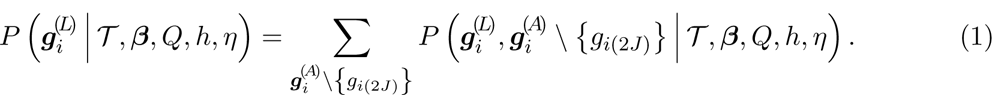

Intuitively, this means that to compute the likelihood of the genotypes of the variant sites at the leaves, we marginalize out the genotypes at the ancestor nodes from the total likelihood. The variables in Equation (1) have the following meaning: *T* is the rooted binary tree topology, whose root, representing a normal cell with diploid genome, has only one child, the most recent common ancestor (MRCA) of all sequenced cells. *T* has *J* sequenced cells as leaves, labelled by {1*, …, J* }, and *J* extinct, ancestor cells as internal nodes, labelled by {*J* + 1*, …*, 2*J* }, where node 2*J* is the root of *T*. The leaves have genotypes ***g***^(^*^L^*^)^_*i*_ = (*g_i_*_1_*, …, g_ij_, …, g_iJ_*)*^T^*, where *g_ij_ ∈ G*, while the internal nodes have genotypes ***g***^(^*^A^*^)^_*i*_ = (*g _i_*_(_*_J_* _+1)_ *, …, g_ij_, …, g _i_*_(2_*_J_* _)_) *^T^*, where *g_ij_ \* {*g _i_*_(2_*_J_* _)_ } *∈ G* and *g_i_*_(2_*_J_* _)_ = 0*/*0. *T* has 2*J −* 1 branches, whose lengths ***β*** *∈* R^2^*^J−^*^1^ represent the expected number of somatic mutations per site. *h* and *η* are the number of rate categories and shape, respectively, of a discrete Gamma distribution with mean equal 1 for modeling among-site substitution rate variation [46]. Hidden random variables *d, T*, ***β****, η* are estimated using MCMC from the posterior of the samples, while the fixed hyperparameter *h* takes value 4 by default.

Given deletion rate *d* (and thus *Q*) and branch length *β*, the seven-by-seven transition probability matrix *R*(*β*) is computed as *R*(*β*) = exp (*Qβ*) [45].

### Model of raw read counts behind DelSIEVE

DelSIEVE’s input data for each cell *j ∈* {1*, …, J* } at each candidate site *i ∈* {1*, …, I*} comes in the form of *D*^(1)^_*ij*_ = (***m****_ij_, c_ij_*), where ***m****_ij_* = {*m_ijk_ | k* = 1, 2, 3} are the read counts of three alternative nucleotides with values in descending order and *c_ij_* is the sequencing coverage (Figure 1a; see Kang *et al.* [32] for explanation of how candidate sites are identified). For acquisition bias correction [47, 48], DelSIEVE also optionally takes raw read count data *D*^(2)^ from *I*^′^ background sites that have a wildtype genotype.

We factorize the probability of observing raw read counts *D_ij_* for cell *j* at site *i* into

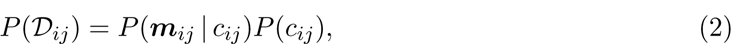

where the former corresponds to the model of nucleotide read counts and the latter to the model of sequencing coverage.

### Model of sequencing coverage

A major, yet often overlooked, challenge in scDNA-seq is the highly uneven sequencing coverage. This happens because the genetic materials are amplified largely unequally during WGA. Similar to SIEVE, we employ a negative binomial distribution to capture the overdispersion existing in the sequencing coverage:

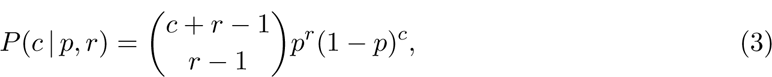

where *p* and *r* are parameters. To improve interpretability, the distribution is reparameterized using mean *µ* and variance *σ*^2^:

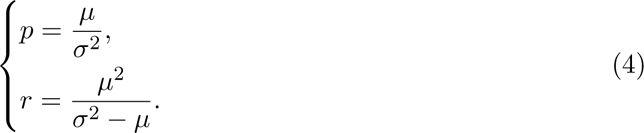

We assume that *µ_ij_*and *σ*^2^_*ij*_ have the same form as in SIEVE, namely

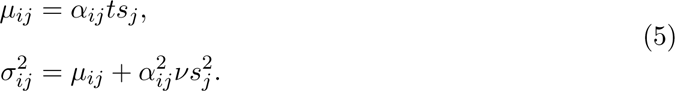

Here, *t* and *ν* are the mean and the variance of allelic coverage, respectively. *α_ij_ ∈* {0, 1, 2} represents the number of sequenced alleles. With the extended genotype state space *G* in the DelSIEVE model, the true number of alleles at a site can either be zero (corresponding to genotype state {-}), one (genotype states {0/-, 1/-}), or two ({0*/*0, 0*/*1, 1*/*1, 1*/*1^′^}). On top of that, the possible occurrence of dropouts during scWGA could also alter the number of observed alleles at a site. Here, we model two types of dropout modes, the loss of one of the two alleles at a site (ADO) or the loss of both alleles (LDO). In ADO simulations and model configuration, only ADO events can occur. In LDO mode, both ADO and LDO may occur. The detailed description of ADO and LDO modes in DelSIEVE is in Additional file 1: Supplementary notes.

In Equation (5) *s_j_* is the size factor of cell *j*, which makes sequencing coverage from different cells comparable, and which is estimated using

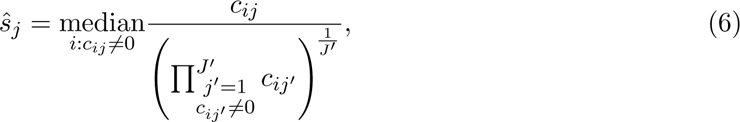

where *J*^′^ is the number of cells with non-zero coverage at a site.

### Model of nucleotide read counts

The occurrence of dropouts could change the number of alleles sequenced for cell *j* at site *i*. As a result, the observed genotype *g*^′^

*∈ G* could be different from the true genotype *g^′^_ij_*. The probability of *g*^′^ is *P* (*g^′^_*ij*_ | g_ij_, α_ij_*), which is defined in Table 1 for the ADO mode and in Table 2 for the LDO mode.

**Table 1:** Definition of the distribution of the observed genotype *g_ij_*^′^ conditional on the true genotype *g_ij_* and number of sequenced alleles *α_ij_* under the ADO mode.

| $g'_{ij}$ | $g_{ij}$ | $\alpha_{ij}$ | $P(g'_{ij} \vert g_{ij}, \alpha_{ij})$ |
| --- | --- | --- | --- |
| 0/0 | 0/0 | 2 | 1 |
| 0/- | 0/0 | 1 | 1 |
| 0/1 | 0/1 | 2 | 1 |
| 0/- | 0/1 | 1 | $\frac{1}{2}$ |
| 1/- | 0/1 | 1 | $\frac{1}{2}$ |
| 1/1 | 1/1 | 2 | 1 |
| 1/- | 1/1 | 1 | 1 |
| 1/1' | 1/1' | 2 | 1 |
| 1/- | 1/1' | 1 | 1 |
| 0/- | 0/- | 1 | 1 |
| - | 0/- | 0 | 1 |
| 1/- | 1/- | 1 | 1 |
| - | 1/- | 0 | 1 |
| - | - | 0 | 1 |
| Others |  |  | 0 |

**Table 2:**
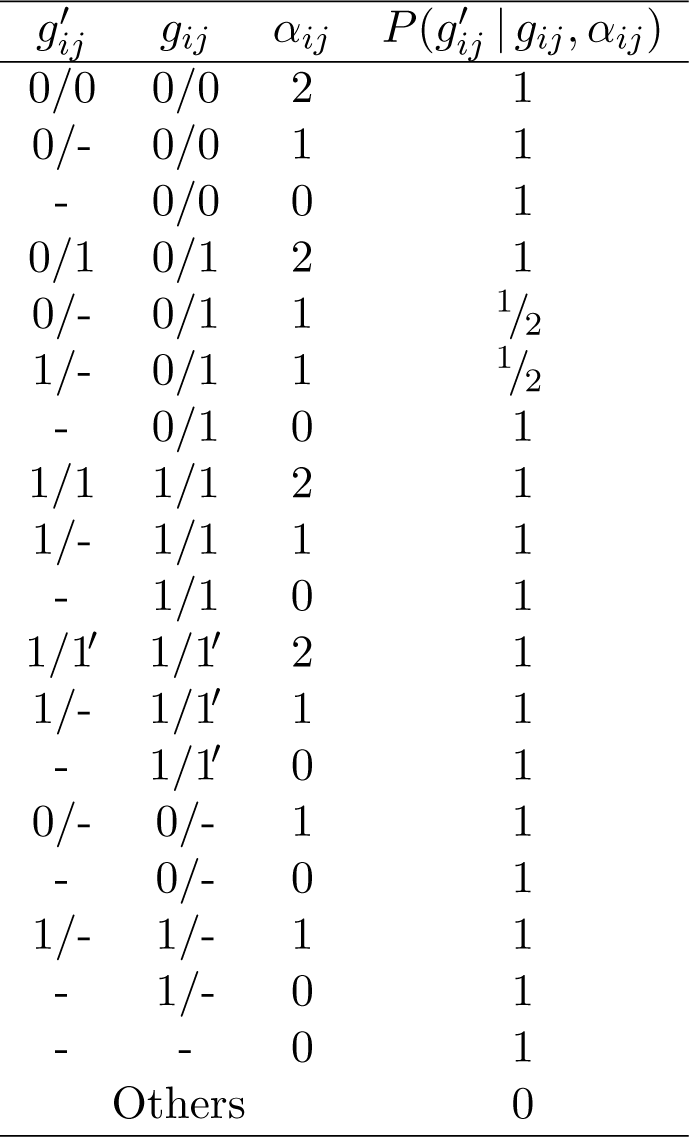
Definition of the distribution of the observed genotype *g^′^_ij_* conditional on the true genotype *g_ij_* and number of sequenced alleles *α_ij_* under the LDO mode.

| $g'_{ij}$ | $g_{ij}$ | $\alpha_{ij}$ | $P(g'_{ij} \vert g_{ij}, \alpha_{ij})$ |
| --- | --- | --- | --- |
| 0/0 | 0/0 | 2 | 1 |
| 0/- | 0/0 | 1 | 1 |
| - | 0/0 | 0 | 1 |
| 0/1 | 0/1 | 2 | 1 |
| 0/- | 0/1 | 1 | $1/2$ |
| 1/- | 0/1 | 1 | $1/2$ |
| - | 0/1 | 0 | 1 |
| 1/1 | 1/1 | 2 | 1 |
| 1/- | 1/1 | 1 | 1 |
| - | 1/1 | 0 | 1 |
| 1/1' | 1/1' | 2 | 1 |
| 1/- | 1/1' | 1 | 1 |
| - | 1/1' | 0 | 1 |
| 0/- | 0/- | 1 | 1 |
| - | 0/- | 0 | 1 |
| 1/- | 1/- | 1 | 1 |
| - | 1/- | 0 | 1 |
| - | - | 0 | 1 |
| Others |  |  | 0 |

When *g^′^_ij_ ∈ G \* {-}, we model ***m****_ij_*, the read counts of three alternative nucleotides, conditional on the sequencing coverage *c_ij_* as a Dirichlet-Multinomial distribution:

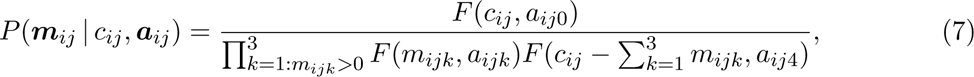

with parameters ***a****_ij_* = {*a_ijk_ | k* = 1*, …*, 4} and 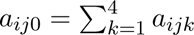. *F* is a function defined as

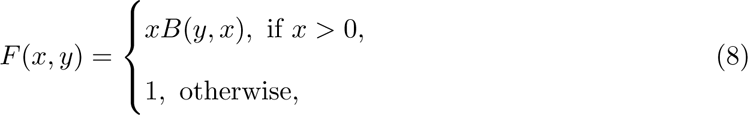

where *B* is the beta function. Note that 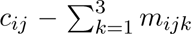 is the read count of the reference nucleotide.

We reparameterize Equation (7) by letting ***a****_ij_* = *w_ij_****f****_ij_*. *w_ij_* captures the overdispersion in the assignment of *c_ij_* read counts among all nucleotides. 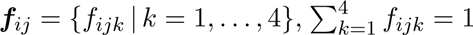 is a vector of expected frequencies of each nucleotide, where the first three elements correspond to the three alternative nucleotides ordered decreasingly according to their read counts, and the last to the reference nucleotide. Depending on *g^′^_ij_*, ***f****_ij_* is given by

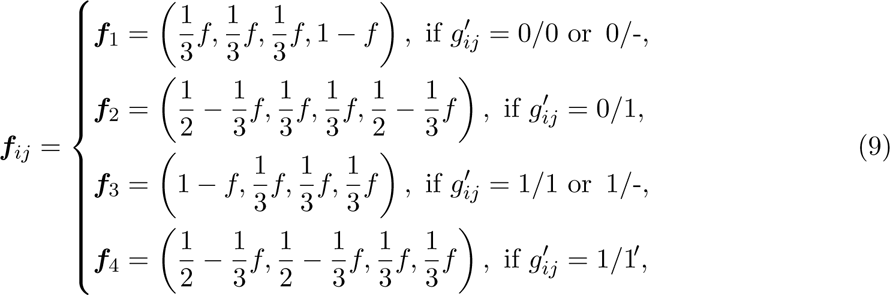

where *f* is the effective sequencing error rate, combining together amplification and sequencing errors.

The parameter *w_ij_*also depends on *g^′^_ij_*, where

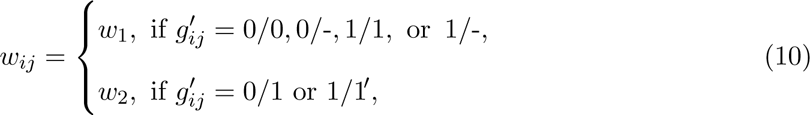

and *w*_1_ is the overdispersion term when *g_ij_*^′^ has only one type of nucelotide, and *w*_2_ is the term when *g^′^_ij_* has different types of nucelotides.

By plugging Equations (9) and (10) into Equation (7), and additionally defining

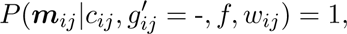

we obtain

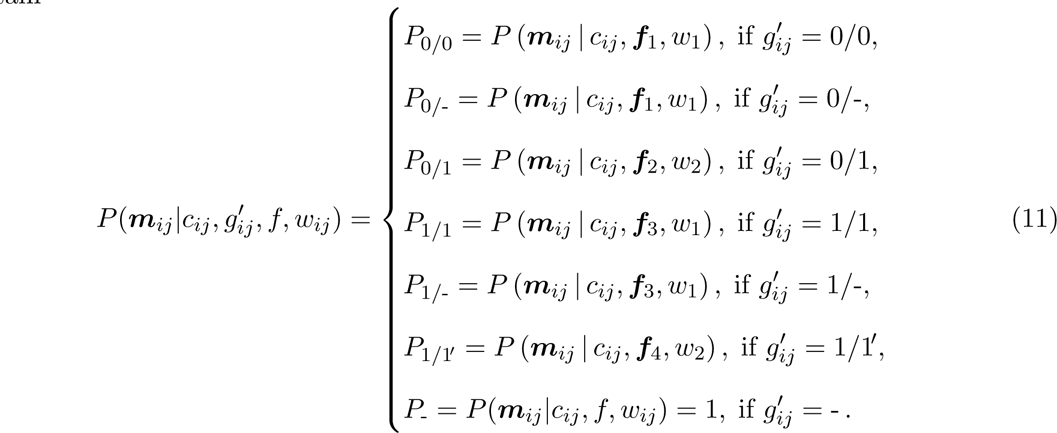

The likelihood, priors and posterior for the DelSIEVE model are defined in Additional file 1: Supplementary notes.

### Mutation event classification

DelSIEVE is able to discern 28 types of genotype transitions, which are classified into 17 types of mutation events (Table 3). Each genotype transition is a possible combination of single point mutations, single back mutations and single deletions. Single point mutations happen when 0 mutates to 1, or 1 and 1^′^ mutate to each other. Single back mutations occur when 1 or 1^′^ mutates to 0. Single deletions happen when an existing allele is lost during evolution, namely 0 or 1 deleted.

**Table 3:** 28 types of genotype transitions that DelSIEVE is able to identify, with their interpretation as mutation events. The genotype transitions correspond to possible changes of genotypes on a branch from the parent node to the child node. If any of these events occurs on independent branches of the phylogenetic tree, it is also considered as a parallel evolution event. The first 12 genotype transitions are also identifiable with SIEVE. LOH in the table represents loss of heterozygosity.

| Genotype transition | Mutation event | Identifiable by |  |
| --- | --- | --- | --- |
|  |  | DelSIEVE | SIEVE |
| 0/0 → 0/1 | Single point mutation | Yes | Yes |
| 0/0 → 1/1 | Coincident homozygous double point mutation | Yes | Yes |
| 0/0 → 1/1' | Coincident heterozygous double point mutation | Yes | Yes |
| 0/1 → 0/0 | Single back mutation | Yes | Yes |
| 1/1 → 0/1 | Single back mutation | Yes | Yes |
| 1/1' → 0/1 | Single back mutation | Yes | Yes |
| 1/1 → 0/0 | Coincident double back mutation | Yes | Yes |
| 1/1' → 0/0 | Coincident double back mutation | Yes | Yes |
| 0/1 → 1/1 | Homozygous single point mutation addition | Yes | Yes |
| 0/1 → 1/1' | Heterozygous single point mutation addition | Yes | Yes |
| 1/1' → 1/1 | Homozygous substitute single point mutation | Yes | Yes |
| 1/1 → 1/1' | Heterozygous substitute single point mutation | Yes | Yes |
| 0/0 → 0/- | Single deletion (not LOH) | Yes | No |
| 1/1 → 1/- | Single deletion (not LOH) | Yes | No |
| 0/1 → 0/- | Single deletion (LOH) | Yes | No |
| 0/1 → 1/- | Single deletion (LOH) | Yes | No |
| 1/1' → 1/- | Single deletion (LOH) | Yes | No |
| 0/0 → 1/- | Coincident deletion and point mutation | Yes | No |
| 1/1 → 0/- | Coincident deletion and back mutation | Yes | No |
| 1/1' → 0/- | Coincident deletion and back mutation | Yes | No |
| 0/- → 1/- | Single deletion point mutation addition | Yes | No |
| 1/- → 0/- | Single deletion back mutation addition | Yes | No |
| 0/- → - | Single deletion addition | Yes | No |
| 1/- → - | Single deletion addition | Yes | No |
| 0/0 → - | Coincident double deletions | Yes | No |
| 0/1 → - | Coincident double deletions | Yes | No |
| 1/1 → - | Coincident double deletions | Yes | No |
| 1/1' → - | Coincident double deletions | Yes | No |

Since DelSIEVE encompasses the genotype state space modeled by SIEVE, it is capable of discerning all genotype transitions that SIEVE can handle, namely the first 12 rows in Table 3 (for detailed explanation see Kang *et al.* [32]). The mutation events that only DelSIEVE is able to discern are explained as follows. Single deletions that happen when one allele is deleted from genotypes in which both alleles originally had different nucleotides result in loss of heterozygosity (LOH) (0*/*1 *→* 0/-, 0*/*1 *→* 1/- and 1*/*1^′^ *→* 1/-). Deletions that take place when one allele is deleted from genotypes in which both alleles originally contained the same nucleotide do not result in LOH (0*/*0 *→* 0/- and 1*/*1 *→* 1/-). The coincident deletion and point mutation type (0*/*0 *→* 1/-) refers to the case when one allele is deleted, and the other is mutated from the wildtype, while the coincident deletion and back mutation (1*/*1 *→* 0/- and 1*/*1^′^ *→* 0/-) happens when one allele is deleted, and the other is mutated back to the reference nucleotide. The single deletion mutation addition (0/- *→* 1/-) takes place when the only allele of the reference-remaining single deletion genotype is mutated to an alternative nucleotide, while the single deletion back mutation addition happens when the mutated allele of the alternative-remaining single deletion genotype is mutated back to the reference nucleotide. The single deletion addition (0/- *→* - and 1/- *→* -) refers to the case when the only allele is deleted of the reference- and alternative-remaining single deletion genotypes. Finally, for the coincident double deletions (0*/*0 *→* -, 0*/*1 *→* -, 1*/*1 *→* - and 1*/*1^′^ *→* -) both of the alleles existing before are deleted.

## Supplementary Materials

### Supplementary Material 1

Supplementary notes, Supplementary Figures. S1-S21 and Tables S1-S4.

## Data availability

We analyzed three published single-cell datasets ([32, 35, 39]). Raw sequencing data for these datasets are available from the BioProject database under accession code PRJNA896550 (CRC28), as well as from the SRA database under accession codes SRA053195 (TNBC16) and SRP067815 (CRC48).

## Code availability

DelSIEVE is implemented in Java and is accessible at https://github.com/szczurek-lab/ DelSIEVE. The simulator is hosted at https://github.com/szczurek-lab/DelSIEVE_simulator, and the reproducible benchmarking framework is available at https://github.com/szczurek-lab/DelSIEVE_benchmark_pipeline. The scripts for generating all figures in this paper are hosted at https://github.com/szczurek-lab/DelSIEVE_analysis. All aforementioned code are freely accessible under a GNU General Public License v3.0 license.

## Acknowledgments

This project has received funding from the European Union’s Horizon 2020 research and innovation programme under the Marie Sk-lodowska-Curie grant agreement No. 766030. E.S. acknowledges the support from the Polish National Science Centre SONATA BIS grant No. 2020/38/E/NZ2/00305. D.P. was supported by the European Research Council (ERC-617457- PHYLOCANCER), the Spanish Ministry of Science and Innovation (PID2019-106247GB-I00), and Xunta de Galicia.

## Author contributions

S.K. and E.S. conceived the DelSIEVE model for which J.K., N.BE. and D.P. provided input and feedback. S.K. implemented the model, benchmarked it and generated the figures. N.BO. and M.V. processed the scDNA-seq datasets. M.M. plotted the copy numbers across the whole genome. S.K. and E.S. wrote the manuscript with critical comments and input from all the co-authors. E.S. supervised the study.

## References

1. Nowell, P. C. The Clonal Evolution of Tumor Cell Populations: Acquired genetic lability permits stepwise selection of variant sublines and underlies tumor progression. Science 194, 23–28 (1976).

2. Hanahan, D. & Weinberg, R. A. The hallmarks of cancer. Cell 100, 57–70 (2000).

3. Hanahan, D. & Weinberg, R. A. Hallmarks of cancer: the next generation. Cell 144, 646– 674 (2011).

4. Vogelstein, B. et al. Cancer genome landscapes. Science 339, 1546–1558 (2013).

5. Hanahan, D. Hallmarks of cancer: new dimensions. Cancer Discovery 12, 31–46 (2022).

6. Greenman, C. et al. Patterns of somatic mutation in human cancer genomes. Nature 446, 153–158 (2007).

7. Stratton, M. R., Campbell, P. J. & Futreal, P. A. The cancer genome. Nature 458, 719–724 (2009).

8. Beroukhim, R. et al. The landscape of somatic copy-number alteration across human cancers. Nature 463, 899–905 (2010).

9. Navin, N. et al. Tumour evolution inferred by single-cell sequencing. Nature 472, 90–94 (2011).

10. Navin, N. E. Cancer genomics: one cell at a time. Genome Biology 15, 1–13 (2014).

11. Navin, N. E. The first five years of single-cell cancer genomics and beyond. Genome Research 25, 1499–1507 (2015).

12. Lähnemann, D., et al. Eleven grand challenges in single-cell data science. Genome Biology 21, 1–35 (2020).

13. Gawad, C., Koh, W. & Quake, S. R. Single-cell genome sequencing: current state of the science. Nature Reviews Genetics 17, 175–188 (2016).

14. Baslan, T. & Hicks, J. Unravelling biology and shifting paradigms in cancer with single-cell sequencing. Nature Reviews Cancer 17, 557–569 (2017).

15. Estévez-Gómez, N. et al. Comparison of single-cell whole-genome amplification strategies. bioRxiv. https://www.biorxiv.org/content/early/2018/10/16/443754 (2018).

16. Mallory, X. F., Edrisi, M., Navin, N. & Nakhleh, L. Methods for copy number aberration detection from single-cell DNA-sequencing data. Genome Biology 21, 1–22 (2020).

17. Laks, E. et al. Clonal decomposition and DNA replication states defined by scaled single-cell genome sequencing. Cell 179, 1207–1221 (2019).

18. Pellegrino, M. et al. High-throughput single-cell DNA sequencing of acute myeloid leukemia tumors with droplet microfluidics. Genome Research 28, 1345–1352 (2018).

19. Zafar, H., Wang, Y., Nakhleh, L., Navin, N. & Chen, K. Monovar: single-nucleotide variant detection in single cells. Nature Methods 13, 505–507 (2016).

20. Dong, X. et al. Accurate identification of single-nucleotide variants in whole-genome-amplified single cells. Nature Methods 14, 491–493 (2017).

21. Bohrson, C. L. et al. Linked-read analysis identifies mutations in single-cell DNA-sequencing data. Nature Genetics 51, 749–754 (2019).

22. Luquette, L. J., Bohrson, C. L., Sherman, M. A. & Park, P. J. Identification of somatic mutations in single cell DNA-seq using a spatial model of allelic imbalance. Nature Communications 10, 1–14 (2019).

23. Yuan, K., Sakoparnig, T., Markowetz, F. & Beerenwinkel, N. BitPhylogeny: a probabilistic framework for reconstructing intra-tumor phylogenies. Genome Biology 16, 1–16 (2015).

24. Ross, E. M. & Markowetz, F. OncoNEM: inferring tumor evolution from single-cell sequencing data. Genome Biology 17, 1–14 (2016).

25. Jahn, K., Kuipers, J. & Beerenwinkel, N. Tree inference for single-cell data. Genome Biology 17, 1–17 (2016).

26. Zafar, H., Tzen, A., Navin, N., Chen, K. & Nakhleh, L. SiFit: inferring tumor trees from single-cell sequencing data under finite-sites models. Genome Biology 18, 1–20 (2017).

27. Malikic, S., Jahn, K., Kuipers, J., Sahinalp, S. C. & Beerenwinkel, N. Integrative inference of subclonal tumour evolution from single-cell and bulk sequencing data. Nature Communications 10, 1–12 (2019).

28. Kozlov, A. M., Darriba, D., Flouri, T., Morel, B. & Stamatakis, A. RAxML-NG: a fast, scalable and user-friendly tool for maximum likelihood phylogenetic inference. Bioinformatics 35, 4453–4455 (2019).

29. Zafar, H., Navin, N., Chen, K. & Nakhleh, L. SiCloneFit: Bayesian inference of population structure, genotype, and phylogeny of tumor clones from single-cell genome sequencing data. Genome Research 29, 1847–1859 (2019).

30. Kozlov, A., Alves, J. M., Stamatakis, A. & Posada, D. CellPhy: accurate and fast probabilistic inference of single-cell phylogenies from scDNA-seq data. Genome Biology 23, 1–30 (2022).

31. Singer, J., Kuipers, J., Jahn, K. & Beerenwinkel, N. Single-cell mutation identification via phylogenetic inference. Nature Communications 9, 5144. 10.1038/ s41467-018-07627-7 (Dec. 2018).

32. Kang, S. et al. SIEVE: joint inference of single-nucleotide variants and cell phylogeny from single-cell DNA sequencing data. Genome Biology 23, 248. 10.1186/ s13059-022-02813-9 (Nov. 2022).

33. Satas, G., Zaccaria, S., Mon, G. & Raphael, B. J. SCARLET: single-cell tumor phylogeny inference with copy-number constrained mutation losses. Cell Systems 10, 323–332 (2020).

34. Kuipers, J., Singer, J. & Beerenwinkel, N. Single-cell mutation calling and phylogenetic tree reconstruction with loss and recurrence. Bioinformatics. btac577. 10.1093/bioinformatics/btac577 (Aug. 2022).

35. Wang, Y. et al. Clonal evolution in breast cancer revealed by single nucleus genome sequencing. Nature 512, 155–160. 10.1038/nature13600 (Aug. 2014).

36. Gao, W.-L., Niu, L., Chen, W.-L., Zhang, Y.-Q. & Huang, W.-H. Integrative analysis of the expression levels and prognostic values for NEK family members in breast cancer. Frontiers in Genetics 13, 798170 (2022).

37. Bersini, S. et al. Nup93 regulates breast tumor growth by modulating cell proliferation and actin cytoskeleton remodeling. Life Science Alliance 3 (2020).

38. Klaus, C. et al. Modulating effects of acyl-CoA synthetase 5-derived mitochondrial Wnt2B palmitoylation on intestinal Wnt activity. World Journal of Gastroenterology: WJG 20, 14855 (2014).

39. Wu, H. et al. Evolution and heterogeneity of non-hereditary colorectal cancer revealed by single-cell exome sequencing. Oncogene 36, 2857–2867 (2017).

40. Raskov, H., Søby, J. H., Troelsen, J., Bojesen, R. D. & Gögenur, I. Driver gene mutations and epigenetics in colorectal cancer. Annals of Surgery 271, 75–85 (2020).

41. D’Andrea, A. D. in The Molecular Basis of Cancer (Fourth Edition) (eds Mendelsohn, J., Gray, J. W., Howley, P. M., Israel, M. A. & Thompson, C. B.) Fourth Edition, 47–66.e2 (W.B. Saunders, Philadelphia, 2015). https://www.sciencedirect.com/science/article/pii/B9781455740666000044.

42. Favero, F. et al. Sequenza: allele-specific copy number and mutation profiles from tumor sequencing data. Annals of Oncology 26, 64–70 (2015).

43. Garvin, T. et al. Interactive analysis and assessment of single-cell copy-number variations. Nature Methods 12, 1058–1060 (2015).

44. Vogelstein, B. & Kinzler, K. W. Cancer genes and the pathways they control. Nature Medicine 10, 789–799 (2004).

45. Felsenstein, J. Inferring phylogenies (Sinauer associates Sunderland, MA, 2004).

46. Yang, Z. Maximum likelihood phylogenetic estimation from DNA sequences with variable rates over sites: approximate methods. Journal of Molecular Evolution 39, 306–314 (1994).

47. Lewis, P. O. A Likelihood Approach to Estimating Phylogeny from Discrete Morphological Character Data. Systematic Biology 50, 913–925. 10.1080/106351501753462876 (Nov. 2001).

48. Leaché, A. D., Banbury, B. L., Felsenstein, J., de Oca, A. n.-M. & Stamatakis, A. Short Tree, Long Tree, Right Tree, Wrong Tree: New Acquisition Bias Corrections for Inferring SNP Phylogenies. Systematic Biology 64, 1032–1047. 10.1093/sysbio/ syv053 (July 2015).

