## Supplementary materials for "DelSIEVE: cell phylogeny model of single nucleotide variants and deletions from single-cell DNA sequencing data"

### Supplementary notes

#### DelSIEVE model: continued

To model ADO mode using DelSIEVE, we introduce the corresponding prior distribution of $\alpha_{ij}$, $P(\alpha_{ij}\,|\,g_{ij},\theta_A)$, as defined in Table S1, where $\theta_A$ denotes the allelic dropout rate under the ADO mode. One should consider the "ADO occurred" column as the value of an additional hidden random variable indicating the occurrence of an ADO, which will be marginalized out in the model. For example, the probability of an ADO when $g_{ij} = 0/\text{-}$ equals ${}^{\theta_A}/_2$, because there is only one allele left to be dropped out. For genotype -, it is certain that ADO has not occurred as there is no allele existing.

To generalize DelSIEVE to model both ADO and LDO, we allow more than one allele to drop out. $P(\alpha_{ij}\,|\,g_{ij},\theta_L)$ is defined in Table S2, where $\theta_L$ represents the allelic dropout rate under the LDO mode. We assume that the ADOs occur to each allele independently. For instance, when $g_{ij} = 0/0$, the probability of $\alpha_{ij} = 0$ is $\theta_L^2$, happening only when both alleles drop out. For genotype 0/-, the sole allele drops out with probability $\theta_L$, resulting in zero sequenced alleles.

#### DelSIEVE likelihood

Combining the statistical phylogenetic model and the model of raw read counts described above, the likelihood of the observed read count data under the DelSIEVE model becomes

$$P\left(\mathcal{D}^{(1)}, \mathcal{D}^{(2)} \,\middle|\, \mathcal{T}, \boldsymbol{\beta}, Q, h, \eta, t, v, \theta, f, w_1, w_2\right). \tag{S.1}$$

To simplify notation, we denote some variables in the statistical phylogenetic model as $\Theta = \{\mathcal{T}, \boldsymbol{\beta}, Q, h, \eta\}$ and some in the model of raw read counts as $\Phi = \{t, v, \theta, f, w_1, w_2\}$. By taking the logarithm, Equation (S.1) becomes

$$\log \mathcal{L}(\Theta, \Phi) = \log \mathcal{L}^{(1)}(\Theta, \Phi) + \log \mathcal{L}^{(2)}(\Theta, \Phi), \tag{S.2}$$

where $\mathcal{L}^{(1)}$ is the tree likelihood corrected for acquisition bias computed for candidate SNV sites in $\mathcal{D}^{(1)}$, while $\mathcal{L}^{(2)}$ is the likelihood computed for background sites in $\mathcal{D}^{(2)}$, referred to as the background likelihood.

Acquisition bias in the context of phylogenetic reconstruction refers to the overestimation of branch lengths (expected number of mutations per site) due to the use of only variable sites [48,

49]. Here, we correct for it following [50]:

$$\log \mathcal{L}^{(1)} = \log P\left(\mathcal{D}^{(1)} \,\middle|\, \Theta, \Phi\right) + I' \log\left(\frac{1}{I}\sum_{i=1}^{I} C_i\right), \tag{S.3}$$

where the first component is the uncorrected log-likelihood for SNV sites, and $C_i$ in the second component is the likelihood of SNV site $i$ being invariant (see below).

To compute $\log P\left(\mathcal{D}^{(1)} \,\middle|\, \Theta, \Phi\right)$ in Equation (S.3), we decompose it according to the probabilistic graphical model in Figure 1b. Assuming independent and identical evolution of each candidate variant site, $\log P\left(\mathcal{D}^{(1)} \,\middle|\, \Theta, \Phi\right)$ is defined as:

$$\begin{aligned}
\log P\left(\mathcal{D}^{(1)} \,\middle|\, \Theta, \Phi\right) &= \sum_{i=1}^{I} \log \sum_{\boldsymbol{g}_i^{(L)}, \boldsymbol{g}_i^{(A)} \setminus \{g_{i(2J)}\}} \Bigg[ P\left(\mathcal{D}_i^{(1)} \,\middle|\, \boldsymbol{g}_i^{(L)}, \Phi\right) \\
&\qquad \times P\left(\boldsymbol{g}_i^{(L)}, \boldsymbol{g}_i^{(A)} \setminus \{g_{i(2J)}\} \,\middle|\, \Theta\right) \Bigg] \\
&= \sum_{i=1}^{I} \log \sum_{\boldsymbol{g}_i^{(L)}, \boldsymbol{g}_i^{(A)} \setminus \{g_{i(2J)}\}} \Bigg[ \prod_{j=1}^{J} P\left(\boldsymbol{m}_{ij}, c_{ij} \,|\, g_{ij}, \Phi\right) \\
&\qquad \times P\left(\boldsymbol{g}_i^{(L)}, \boldsymbol{g}_i^{(A)} \setminus \{g_{i(2J)}\} \,\middle|\, \Theta\right) \Bigg] \\
&= \sum_{i=1}^{I} \sum_{j=1}^{J} \log \sum_{\boldsymbol{g}_i^{(L)}, \boldsymbol{g}_i^{(A)} \setminus \{g_{i(2J)}\}} \Bigg[ P\left(\boldsymbol{m}_{ij}, c_{ij} \,|\, g_{ij}, \Phi\right) \\
&\qquad \times P\left(\boldsymbol{g}_i^{(L)}, \boldsymbol{g}_i^{(A)} \setminus \{g_{i(2J)}\} \,\middle|\, \Theta\right) \Bigg],
\end{aligned} \tag{S.4}$$

where $P(\boldsymbol{m}_{ij}, c_{ij} \,|\, g_{ij}, \Phi)$, representing the model of raw read counts, is similarly decomposed into

$$\begin{aligned}
P\left(\boldsymbol{m}_{ij}, c_{ij} \,|\, g_{ij}, \Phi\right) &= P\left(\boldsymbol{m}_{ij}, c_{ij} \,|\, g_{ij}, f, w_{ij}, t, v, \theta\right) \\
&= \sum_{\alpha_{ij}, g'_{ij}} P\left(\boldsymbol{m}_{ij}, c_{ij}, \alpha_{ij}, g'_{ij} \,|\, g_{ij}, f, w_{ij}, t, v, \theta\right) \\
&= \sum_{\alpha_{ij}, g'_{ij}} \Big[ P\left(\boldsymbol{m}_{ij} \,|\, c_{ij}, g'_{ij}, f, w_{ij}\right) P\left(g'_{ij} \,|\, g_{ij}, \alpha_{ij}\right) \\
&\qquad \times P\left(c_{ij} \,|\, \alpha_{ij}, t, v\right) P\left(\alpha_{ij} \,|\, g_{ij}, \theta\right) \Big].
\end{aligned} \tag{S.5}$$

$P(c_{ij} \,|\, \alpha_{ij}, t, v)$ in Equation (S.5) is defined through Equations (3)-(5), and $P(\boldsymbol{m}_{ij} \,|\, c_{ij}, g'_{ij}, f, w_{ij})$ is defined in Equation (11). Under the ADO mode, $P(\alpha_{ij} \,|\, g_{ij}, \theta)$ and $P(g'_{ij} \,|\, g_{ij}, \alpha_{ij})$ are defined as shown in Table S1 and Table 1, respectively, while under the LDO mode in Table S2 and Table 2, respectively. As a result, Equation (S.5) takes distinct forms under different dropout modes.

For the ADO mode, Equation (S.5) is further represented as

$$P\left(\boldsymbol{m}_{ij}, c_{ij} \,|\, g_{ij}, \Phi\right) = \begin{cases} P_{0/0} \cdot P(c_{ij} \,|\, \alpha_{ij} = 2, t, v) \cdot (1-\theta) \\ \qquad + P_{0/\text{-}} \cdot P(c_{ij} \,|\, \alpha_{ij} = 1, t, v) \cdot \theta, \text{ if } g_{ij} = 0/0, \\ P_{0/1} \cdot P(c_{ij} \,|\, \alpha_{ij} = 2, t, v) \cdot (1-\theta) \\ \qquad + \frac{1}{2}(P_{0/\text{-}} + P_{1/\text{-}}) \cdot P(c_{ij}|\alpha_{ij} = 1, t, v) \cdot \theta, \text{ if } g_{ij} = 0/1, \\ P_{1/1} \cdot P(c_{ij} \,|\, \alpha_{ij} = 2, t, v) \cdot (1-\theta) \\ \qquad + P_{1/\text{-}} \cdot P(c_{ij} \,|\, \alpha_{ij} = 1, t, v) \cdot \theta, \text{ if } g_{ij} = 1/1, \\ P_{1/1'} \cdot P(c_{ij} \,|\, \alpha_{ij} = 2, t, v) \cdot (1-\theta) \\ \qquad + P_{1/\text{-}} \cdot P(c_{ij} \,|\, \alpha_{ij} = 1, t, v) \cdot \theta, \text{ if } g_{ij} = 1/1', \\ P_{0/\text{-}} \cdot P(c_{ij} \,|\, \alpha_{ij} = 1, t, v) \cdot (1 - \frac{\theta}{2}) \\ \qquad + P_{\text{-}} \cdot P(c_{ij} \,|\, \alpha_{ij} = 0, t, v) \cdot \frac{\theta}{2}, \text{ if } g_{ij} = 0/\text{-}, \\ P_{1/\text{-}} \cdot P(c_{ij} \,|\, \alpha_{ij} = 1, t, v) \cdot (1 - \frac{\theta}{2}) \\ \qquad + P_{\text{-}} \cdot P(c_{ij} \,|\, \alpha_{ij} = 0, t, v) \cdot \frac{\theta}{2}, \text{ if } g_{ij} = 1/\text{-}, \\ P_{\text{-}} \cdot P(c_{ij} \,|\, \alpha_{ij} = 0, t, v), \text{ if } g_{ij} = \text{-}\,. \end{cases} \tag{S.6}$$

For the LDO mode, Equation (S.5) is

$$P\left(\boldsymbol{m}_{ij}, c_{ij} \,|\, g_{ij}, \Phi\right) = \begin{cases} P_{0/0} \cdot P(c_{ij} \,|\, \alpha_{ij} = 2, t, v) \cdot (1-\theta)^2 \\ \qquad + P_{0/\text{-}} \cdot P(c_{ij} \,|\, \alpha_{ij} = 1, t, v) \cdot 2 \cdot \theta \cdot (1-\theta) \\ \qquad + P_{\text{-}} \cdot P(c_{ij} \,|\, \alpha_{ij} = 0, t, v) \cdot \theta^2, \text{ if } g_{ij} = 0/0, \\ P_{0/1} \cdot P(c_{ij} \,|\, \alpha_{ij} = 2, t, v) \cdot (1-\theta)^2 \\ \qquad + (P_{0/\text{-}} + P_{1/\text{-}}) \cdot P(c_{ij} | \alpha_{ij} = 1, t, v) \cdot \theta \cdot (1-\theta) \\ \qquad + P_{\text{-}} \cdot P(c_{ij} \,|\, \alpha_{ij} = 0, t, v) \cdot \theta^2, \text{ if } g_{ij} = 0/1, \\ P_{1/1} \cdot P(c_{ij} \,|\, \alpha_{ij} = 2, t, v) \cdot (1-\theta)^2 \\ \qquad + P_{1/\text{-}} \cdot P(c_{ij} \,|\, \alpha_{ij} = 1, t, v) \cdot 2 \cdot \theta \cdot (1-\theta) \\ \qquad + P_{\text{-}} \cdot P(c_{ij} \,|\, \alpha_{ij} = 0, t, v) \cdot \theta^2, \text{ if } g_{ij} = 1/1, \\ P_{1/1'} \cdot P(c_{ij} \,|\, \alpha_{ij} = 2, t, v) \cdot (1-\theta)^2 \\ \qquad + P_{1/\text{-}} \cdot P(c_{ij} \,|\, \alpha_{ij} = 1, t, v) \cdot 2 \cdot \theta \cdot (1-\theta) \\ \qquad + P_{\text{-}} \cdot P(c_{ij} \,|\, \alpha_{ij} = 0, t, v) \cdot \theta^2, \text{ if } g_{ij} = 1/1', \\ P_{0/\text{-}} \cdot P(c_{ij} \,|\, \alpha_{ij} = 1, t, v) \cdot (1-\theta) \\ \qquad + P_{\text{-}} \cdot P(c_{ij} \,|\, \alpha_{ij} = 0, t, v) \cdot \theta, \text{ if } g_{ij} = 0/\text{-}, \\ P_{1/\text{-}} \cdot P(c_{ij} \,|\, \alpha_{ij} = 1, t, v) \cdot (1-\theta) \\ \qquad + P_{\text{-}} \cdot P(c_{ij} \,|\, \alpha_{ij} = 0, t, v) \cdot \theta, \text{ if } g_{ij} = 1/\text{-}, \\ P_{\text{-}} \cdot P(c_{ij} \,|\, \alpha_{ij} = 0, t, v), \text{ if } g_{ij} = \text{-}\,. \end{cases} \tag{S.7}$$

Equation (S.4) is computed efficiently using Felsenstein's pruning algorithm [51]. For $I$ candidate SNV sites, $J$ cells and $K$ genotype states in $G$ (for DelSIEVE $K = 7$), the time complexity of the Felsenstein's pruning algorithm is $\mathcal{O}(IJK^2)$.

Since in the second component of Equation (S.3), $C_i$ corresponds to the likelihood of candidate SNV site $i$ being invariant, it is computed as the joint probability of $\mathcal{D}_i$ and $\boldsymbol{g}_i^{(L)} = 0/0$:

$$\begin{aligned} C_i &= P\left(\mathcal{D}_i^{(1)}, \boldsymbol{g}_i^{(L)} = 0/0 \,\middle|\, \Theta, \Phi\right) \\ &= P\left(\mathcal{D}_i^{(1)} \,\middle|\, \boldsymbol{g}_i^{(L)} = 0/0, \Phi\right) \sum_{\boldsymbol{g}_i^{(A)} \setminus \{g_{i(2J)}\}} P\left(\boldsymbol{g}_i^{(L)} = 0/0, \boldsymbol{g}_i^{(A)} \setminus \{g_{i(2J)}\} \,\middle|\, \Theta\right) \\ &= \prod_{j=1}^{J} P\left(\boldsymbol{m}_{ij}, c_{ij} \,|\, g_{ij} = 0/0, \Phi\right) \sum_{\boldsymbol{g}_i^{(A)} \setminus \{g_{i(2J)}\}} P\left(\boldsymbol{g}_i^{(L)} = 0/0, \boldsymbol{g}_i^{(A)} \setminus \{g_{i(2J)}\} \,\middle|\, \Theta\right), \end{aligned} \tag{S.8}$$

which is computed similarly to Equation (S.4), but with $g_{ij}$ for $j = 1, \dots J$ fixed to $0/0$. In fact, $C_i$ and $\log P\left(\mathcal{D}_i^{(1)} \,\middle|\, \Theta, \Phi\right)$ are computed simultaneously in the implementation for optimized efficiency.

To efficiently compute $\log \mathcal{L}^{(2)}$, the background likelihood in Equation (S.2), we make several simplifications similar to SIEVE. Specifically, we assume that each cell at each background site has the wildtype genotype and is sequenced without dropouts and with at least one read per allele. We also assume that $P(c_{ij} \,|\, \alpha_{ij}, t, v) = 1$ and $P\left(\boldsymbol{g}_i^{(L)} = 0/0, \boldsymbol{g}_i^{(A)} \setminus \{g_{i(2J)}\} \,\middle|\, \Theta\right) = 1$, thereby ignoring the model of sequencing coverage and the tree log-likelihood for the background sites $i$ for $i = 1, \dots I'$. With an alternative form of the Dirichlet-multinomial distribution, $\log \mathcal{L}^{(2)}$ is approximately and efficiently computed by

$$
\begin{aligned}
\log \mathcal{L}^{(2)}(f, w_1) &= \sum_{i=1}^{I'} \sum_{j=1}^{J} \log P_{0/0} \\
&= \sum_{i=1}^{I'} \sum_{j=1}^{J} \log \left[ \frac{\Gamma(w_1)\Gamma(c_{ij}+1)}{\Gamma(c_{ij}+w_1)} \prod_{k=1}^{3} \frac{\Gamma\left(m_{ijk} + \frac{1}{3} f w_1\right)}{\Gamma\left(\frac{1}{3} f w_1\right)\Gamma(m_{ijk}+1)} \right. \\
&\qquad\qquad \left. \times \frac{\Gamma\left(c_{ij} - \sum_{k=1}^{3} m_{ijk} + (1-f)w_1\right)}{\Gamma((1-f)w_1)\Gamma\left(c_{ij} - \sum_{k=1}^{3} m_{ijk} + 1\right)} \right] \\
&= I'J \left[ \log \Gamma(w_1) - 3 \log \Gamma\left(\frac{1}{3} f w_1\right) - \log \Gamma((1-f)w_1) \right] \\
&\quad + \sum_{c=1}^{\max(c_{ij})} N_c (\log \Gamma(c+1) - \log \Gamma(c+w_1)) \\
&\quad + \sum_{k=1}^{3} \sum_{m_k=1}^{\max(m_{ijk})} N_{m_k} \left( \log \Gamma\left(m_k + \frac{1}{3} f w_1\right) - \log \Gamma(m_k+1) \right) \\
&\quad + \sum_{c - \sum_{k=1}^{3} m_k = 1}^{\max(c_{ij} - \sum_{k=1}^{3} m_{ijk})} N_{c - \sum_{k=1}^{3} m_k} \left( \log \Gamma\left(c - \sum_{k=1}^{3} m_k + (1-f)w_1\right) \right. \\
&\qquad\qquad \left. - \log \Gamma\left(c - \sum_{k=1}^{3} m_k + 1\right) \right),
\end{aligned}
\tag{S.9}
$$

where $P_{0/0}$ is defined in Equation (11). Across $I'$ background sites and $J$ cells, $N_c$, $N_{m_k}$ for $k = 1, 2, 3$, and $N_{c - \sum_{k=1}^{3} m_k}$ represent the unique occurrences of sequencing coverage $c$, of alternative nucleotide read counts $m_k$ for $k = 1, 2, 3$, and of reference nucleotide read counts $c - \sum_{k=1}^{3} m_k$, respectively. Some terms, namely $\log \Gamma(c+1)$, $-\log \Gamma(m_k+1)$ for $k = 1, 2, 3$, and $-\log \Gamma\left(c - \sum_{k=1}^{3} m_k + 1\right)$, are constants, and thus they are not updated in the MCMC iterations.

The time complexity of Equation (S.9) is $\mathcal{O}(c)$, where $c$ is the number of unique values in the set of values representing sequencing coverage and read counts for all four nucleotides across

$I'$ background sites and $J$ cells. Since generally $IJK^2 \gg c$, the overall time complexity of model likelihood is $\mathcal{O}(IJK^2)$. It is worth noting that given $I$ candidate variant sites and $J$ cells, the time complexity of DelSIEVE is around 1.8 times greater than that of SIEVE due to the expanded genotype state space.

### Priors

Similar to SIEVE, we use prior distributions predefined and implemented in BEAST 2 for some of the hidden random variables in the DelSIEVE model. For the cell phylogeny given by $\mathcal{T}$ and $\boldsymbol{\beta}$, we set a prior following the Kingman coalescent process with an exponentially growing population, denoted

$$P(\mathcal{T}, \boldsymbol{\beta} \,|\, M, e), \tag{S.10}$$

where $M$ and $e$ are hidden random variables, representing the scaled population size and the exponential growth rate, respectively. The analytical form of Equation (S.10) is defined at length in [52].

The default assumed distribution for $M$ in BEAST 2 is

$$P(M \,|\, \delta) = \frac{1}{\delta}, \tag{S.11}$$

where $\delta$ is the current proposed value of $M$.

As for $e$, the default prior is

$$e \,|\, \lambda, \epsilon \sim \text{Laplace}(\lambda, \epsilon), \tag{S.12}$$

where the default values of the fixed parameters are mean $\lambda = 10^{-3}$ and scale $\epsilon = 30.7$.

For $\eta$ in Equation (1), an exponential prior distribution is chosen:

$$\eta \,|\, \gamma \sim \exp(\gamma), \tag{S.13}$$

where $\gamma = 1$.

For the relative deletion rate $d$, a uniform prior distribution is used:

$$d \,|\, \varphi \sim \text{Uniform}(0, \varphi), \tag{S.14}$$

where $\varphi = 1$.

For the hidden random variables in the model of sequencing coverage in Equations (3)-(5), a non-informative prior is set for $t$:

$$t \,|\, \rho \sim \text{Uniform}(0, \rho), \tag{S.15}$$

where $\rho = 1000$, while the prior for $v$ is

$$v \,|\, \zeta \sim \exp(\zeta), \tag{S.16}$$

where $\zeta = 25$.

For the allelic dropout rate $\theta_A$ defined under the ADO (Table S1) or $\theta_L$ under the LDO mode (Table S2), we use a non-informative prior:

$$\theta \,|\, u \sim \text{Uniform}(0, u), \tag{S.17}$$

where $u = 1$.

Regarding the hidden random variables in the model of nucleotide read counts in Equations (7), (9) and (10), an exponential prior is set for $f$:

$$f \,|\, \tau \sim \exp(\tau), \tag{S.18}$$

where $\tau = 0.025$, and a log normal prior for both $w_1$ and $w_2$:

$$\begin{aligned} w_1 \,|\, \xi_1, \psi_1 &\sim \text{Log-Normal}(\xi_1, \psi_1), \\ w_2 \,|\, \xi_2, \psi_2 &\sim \text{Log-Normal}(\xi_2, \psi_2), \end{aligned} \tag{S.19}$$

where we choose for $w_1$ the log-transformed mean $\xi_1 = 3.9$ (150 for untransformed) and the standard deviation $\psi_1 = 1.5$, and for $w_2$ the log-transformed mean $\xi_2 = 0.9$ (10 for untransformed) and the standard deviation $\psi_2 = 1.7$. The mean is log-transformed using

$$\xi_{\text{transformed}} = \log(\xi_{\text{untransformed}}) - \frac{\psi^2}{2}.$$

These values of the fixed parameters in Equation (S.19) are chosen to cover a wide range of possible values for $w_1$ and $w_2$.

### Posterior and MCMC

The posterior distribution of the hidden random variables is:

$$
\begin{aligned}
& P\left(\mathcal{T}, \boldsymbol{\beta}, M, e, \eta, d, t, v, \theta, f, w_1, w_2 \,\middle|\, \mathcal{D}^{(1)}, \mathcal{D}^{(2)}\right) \\
=& \frac{1}{Z} P\left(\mathcal{D}^{(1)}, \mathcal{D}^{(2)} \,\middle|\, \mathcal{T}, \boldsymbol{\beta}, Q, \eta, t, v, \theta, f, w_1, w_2\right) \\
& \times P(\mathcal{T}, \boldsymbol{\beta} \,|\, M, e) P(M \,|\, \delta) P(e \,|\, \lambda, \epsilon) \\
& \times P(\eta \,|\, \gamma) P(Q \,|\, d) P(d \,|\, \varphi) \\
& \times P(t \,|\, \rho) P(v \,|\, \zeta) P(\theta \,|\, u) P(f \,|\, \tau) \\
& \times P(w_1 \,|\, \xi_1, \psi_1) P(w_2 \,|\, \xi_2, \psi_2),
\end{aligned}
\tag{S.20}
$$

where $Z = P(\mathcal{D}^{(1)}, \mathcal{D}^{(2)})$ is a normalization constant, and the likelihood of the model and priors for hidden random variables are defined in Section DelSIEVE likelihood and Section Priors, respectively. To simplify the notation, we denote the hidden random variables in Equation (S.20) as $\Lambda = \{\mathcal{T}, \boldsymbol{\beta}, M, e, \eta, d, t, v, \theta, f, w_1, w_2\}$.

Since $Z$ in Equation (S.20) is intractable to calculate, we leverage BEAST2's MCMC algorithm with Metropolis-Hastings kernel to sample from the posterior distribution. In this algorithm, a new state of the hidden random variables $\Lambda^*$ is proposed based on its current state $\Lambda$ following a proposal distribution $q(\Lambda^* \,|\, \Lambda)$. $q(\Lambda^* \,|\, \Lambda)$ is designed to ensure the reversibility and ergodicity of the underlying Markov chain. For DelSIEVE, in each iteration, a new state of a randomly selected hidden variable is accepted with probability

$$
\min \left\{1, \frac{P\left(\Lambda^* \,\middle|\, \mathcal{D}^{(1)}, \mathcal{D}^{(2)}\right) q(\Lambda \,|\, \Lambda^*)}{P\left(\Lambda \,\middle|\, \mathcal{D}^{(1)}, \mathcal{D}^{(2)}\right) q(\Lambda^* \,|\, \Lambda)}\right\}. \tag{S.21}
$$

We employ exactly the same proposal distributions as in SIEVE. Briefly, regarding the branch lengths of the tree, the heights of the internal nodes are adjusted. For the tree topology, we use multiple moves, including subtree swapping, Wilson-Balding, and subtree sliding, where the last two moves also change branch lengths as a side effect. With respect to unknown parameters, scaling and random Gaussian walks are used. For detailed description of the aforementioned moves, refer to Drummond et al. [52] and Kang et al. [32].

To achieve more accurate parameter and tree estimates, DelSIEVE employs a two-stage sampling strategy similar to SIEVE.

### Variant calling, dropout calling, and maximum likelihood gene annotation

In the computation of model likelihood using Equations (S.4) and (S.5), we marginalize out some hidden random variables: $\boldsymbol{g}_i^{(L)}$, $\boldsymbol{g}_i^{(A)}$, $g'_{ij}$ and $\alpha_{ij}$. Hence, the direct results from the MCMC sampling process are the posterior distributions of the cell phylogeny and other unknown hidden random variables. We obtain the estimates of those marginalized hidden random variables in a post-processing step. Specifically, we use the max-sum algorithm [53], by fixing the maximum clade credibility tree [54] and parameters estimated from the MCMC posterior samples using TreeAnnotator and Tracer [55], respectively. As a result, genotypes, dropout states, as well as the locations of mutated genes on the inferred cell phylogeny are determined by identifying the maximum likelihood states of $\boldsymbol{g}_i^{(L)}$, $g'_{ij}$ and $\alpha_{ij}$, as well as $\boldsymbol{g}_i^{(A)}$, respectively.

### ScDNA-seq data simulator

We generated simulated data by modifying the simulator we had used in Kang *et al.* [32]. The simulator builds upon a previous simulation method, Cellcoal [56]. The first change we made was to expand the rate matrix, according to which each genomic site evolved along the tree (Table S3). The rate matrix contains 14 genotypes encoded with nucleotides, allowing for point mutations, back point mutations, and deletions. It has one parameter, the deletion rate, which is measured relatively to the mutation rate. Another change was that we implemented the LDO mode to allow more than one dropout to occur at each site for each cell. The simulator takes the same input configuration as the simulator in Kang *et al.* [32] does.

The simulation process was similar to that in Kang *et al.* [32]. Briefly, with a given number of cells, a binary cell lineage tree was first simulated following a neutral coalescent process with exponential growth under a strict molecular clock, where the mutation rate is constant. For a given number of genomic sites, each site was initialized by randomly selecting one of four nucleotides to have a reference genotype. Next, with a given mutation rate and a relative deletion rate, each site was evolved independently along the tree following the rate matrix defined in Table S3. A genomic site was considered as a true SNV site if at least one cell had a genotype that was not wildtype. Dropouts were then added on top of the simulated genotypes under either ADO or LDO mode, as long as there were non-deleted alleles. We recorded the true dropout states for all cells at the true SNV sites. Size factors in Equation (6) were generated from a normal distribution with the mean = 1.2 and the variance = 0.2. The sequencing coverage was simulated using a negative binomial distribution following Equations (3)-(5). The read counts of each nucleotide were then generated following a multinomial distribution.

### Simulation design

We designed a series of simulations to benchmark the performance of DelSIEVE. We reused and modified the benchmarking framework we used for SIEVE.

We assumed that 40 tumor cells were sampled from an exponentially growing population, whose growth rate and effective population size are $10^{-4}$ and $10^4$, respectively. We used the same mutation rates as in the SIEVE benchmark, namely $10^{-6}$, $8 \times 10^{-6}$ and $3 \times 10^{-5}$. We selected two relative deletion rates: 0.1 and 0.25.

For each mutation rate, we simulated a number of genomic sites such that the preselection procedure would produce a certain amount of candidate variant sites, and additionally a high enough number of background sites so that they provide signal for branch length correction. At the same time, we kept the overall number of sites limited for computational efficiency reasons. Specifically, for a mutation rate of $10^{-6}$, we evolved $10^4$ genomic sites to have around $400 \sim 700$ candidate variant sites. For this simulated data, the resulting ratio of background to SNV sites was around 5. For a mutation rate of $8 \times 10^{-6}$, $10^4$ genomic sites were chosen to have around $4 \times 10^3$ background sites. For mutation rate $3 \times 10^{-5}$, $1.2 \times 10^5$ genomic sites were chosen to have at least $2.5 \times 10^3$ background sites. For the higher mutation rates of $8 \times 10^{-6}$ and $3 \times 10^{-5}$, the chosen numbers of genomic sites resulted in $> 5 \times 10^3$ and $> 1.1 \times 10^5$ true SNV sites, respectively. Next, the obtained genomic sites were subsetted to ensure that the number of true SNV sites in the final simulated data for different mutation rates were within the same range, and that for the higher mutation rates $8 \times 10^{-6}$ and $3 \times 10^{-5}$ the ratio between the number of background sites and the true SNV sites was kept similar to the data for the lowest mutation rate (at least 5). To this end, we first computed a targeted number of true SNV sites for each simulated dataset $n_{\text{target}}$ using

$$n_{\text{target}} = \min(700, \frac{n'}{5}),$$

where $n'$ is the number of background sites. Next, we randomly selected $n_{\text{target}}$ sites out of the true SNV sites. Together with the $n'$ background sites, the selected $n_{\text{target}}$ true SNV sites formed the new simulated data.

We considered both ADO and LDO mode. The allelic dropout rate for the former was $\theta_A = 0.3$ , and for the latter $\theta_L = 0.163$ (both are the default setups in Cellcoal [56]).

We had different combinations of $t$ and $v$ in Equations (3)-(5) for various coverage qualities. For simulated data referred to as high coverage quality, we used high mean ($t = 20$) and low variance ($v = 2$) of allelic coverage. For medium coverage quality data, we used high mean ($t = 20$) and medium variance ($v = 10$). For low coverage quality data, we fixed low mean ($t = 5$) and high variance ($v = 20$).

Other parameters were fixed when simulating the data. We set $w_1$ and $w_2$ in Equation (10) to 100 and 2.5, respectively. Moreover, we set both the amplification and sequencing error rate to $10^{-3}$, and thus the effective sequencing error rate in Equation (9) was $f \approx 2 \times 10^{-3}$.

Overall, we designed 36 simulation scenarios, each repeated 10 times.

Furthermore, for each of those genotypes related to deletions, we filtered out results if the proportion of simulated ground truth was less than 0.1%. We also excluded results with a mutation rate of $10^{-6}$ as too few deletions were generated (less than 0.3%, 0.7% and 0.005% for alternative-left single deletion, reference-left single deletion and double deletions, respectively). For the same reason, results were also excluded for double deletions with a mutation rate of $8 \times 10^{-6}$ and for double mutant genotypes with a mutation rate of $10^{-6}$, both of which had less than 0.2% corresponding genotypes generated.

### Variant calling and phylogenetic accuracy

For assessing the results of variant and dropout calling, standard performance measures such as precision, recall, F1 score, and false positive rate (FPR) were used. These measures were computed for DelSIEVE and SIEVE in the task of calling deletions as well as dropouts, and computed for Monovar in addition to DelSIEVE and SIEVE in the task of calling single and double mutant. Moverover, in order to compare to SCIPhIN, these measures were also computed for DelSIEVE, SIEVE and Monovar with respect to the occurrence of deletions and / or single and double mutant (see Section Configurations of methods in Supplementary notes).

Both DelSIEVE and SCIPhIN identify deletions at preselected candidate sites. Hence, we subsetted the true deletions to those at the candidate variant sites when computing the metrics.

To assess the accuracy of cell phylogeny reconstruction, we used the same measurements as in Kang *et al.* [32], namely the BS distance [57] for both the tree topology and branch lengths, as well as the normalized RF distance [58] for the tree topology only (see Kang *et al.* [32]). For DelSIEVE, SIEVE and SiFit, we computed both the BS and the normalized RF distance in the rooted tree mode. For SCIPhIN, we only computed the normalized RF distance as it only infers a rooted tree without branch lengths. We used the R package phangorn to compute the BS and normalized RF distances [59].

### Configurations of methods

For Monovar (commit 68fbb68), we used the true values of $\theta$ and $f$ as priors for the false negative and false positive rates, and default values for other options.

For SCIPhIN (commit 27e5ca6), we gave it the true value of $f$ to avoid estimating its mean error rate (option "wildMean"), and ran it with $10^6$ iterations with zygosity learned (option "lz" set to 1). We also set the penalty of computing the loss (option "llp") and parallel score (option "lpp") to 30. The command line was as follows:

```
sciphin -l 1000000 --lz 1 --ll 1 --lp 1 --llp 30 --lpp 30 --ese 0 \
--wildMean 0.002
```

Note that SCIPhIN does not in its output differentiate deletions from SNVs; that is, it reports both SNVs and deletions simply as mutations.

To run SiFit (commit 9dc3774), we fed the required data with variants called by Monovar using a ternary matrix. We used the true values of $\theta$ and $f$ as the prior for false negative rate and the estimated false positive rate, respectively. We ran it with $2 \times 10^5$ iterations.

For SIEVE, originally it only supported ADO mode. To allow comparison to DelSIEVE in LDO mode, we implemented LDO for SIEVE as well.

We enforced a strict molecular clock model for DelSIEVE and SIEVE, both of which were run for $2 \times 10^6$ and $1.5 \times 10^6$ iterations for the first and the second sampling stages, respectively. The deletion rate was inferred in the second sampling stage as it is related to the branch lengths of the cell phylogeny. Both DelSIEVE and SIEVE were configured to match the dropout mode (ADO or LDO) employed during the simulation process. Additionally, it is necessary to evaluate the performance of DelSIEVE and SIEVE when their configured dropout modes are distinct from that used to simulate a specific dataset. To this end, we selected the extremest simulation scenarios for both the dropout modes (see Section Simulation design in Supplementary notes), namely the lowest and highest mutation rates ($10^{-6}$ and $3 \times 10^{-5}$, respectively), the lowest and highest coverage qualities ($t = 20$, $v = 2$ and $t = 5$, $v = 20$, respectively), and the highest relative deletion rate (0.25).

On the real datasets, we used a log-normal relaxed molecular clock model to account for branch-wise substitution rate variation for DelSIEVE. To obtain better mixed Markov chains, we used an optimized relaxed clock model [60] rather than the default one in BEAST 2. We increased the number of iterations for both stages to $4\times10^6$ and $3.5\times10^6$, respectively. Both the deletion rate and parameters introduced by the relaxed molecular clock model were explored in the second sampling stage. As the simulation results showed that the performance of DelSIEVE is not sensitive to the configured dropout mode, we applied DelSIEVE in ADO mode to the real datasets.

To run Ginkgo on the real datasets, sample bam files were first converted to bed files using `bamtobed` function from bedtools (v2.28.0). With the bed files as input, we ran Ginkgo with default settings, where hg19 reference genome, 1,000,000 bin size and bwa were used as the aligner, and euclidian distance as the clustering option.

To run Sequenza on the real datasets, we used the `bam2seqz` command in the sequenza-utils package to convert bam files for the corresponding bulk samples to the Sequenza file format, which was subsequently binned with the seqz_binning command, using a window size of 50. With this file as input, we used the `sequenza.fit` command from Sequenza v3.0.0 to estimate the ploidy.

SNVs were annotated using Annovar (version 2020 Jun. 08) [61]. The cell phylogeny was plotted in R (version 4.2.3) [62] using ggtree [63], and the genotype heatmap was plotted using ComplexHeatmap [64]. Besides, the comparison of sequencing coverages reported by DelSIEVE and Sequenza was performed and plotted using ggstatsplot [65].

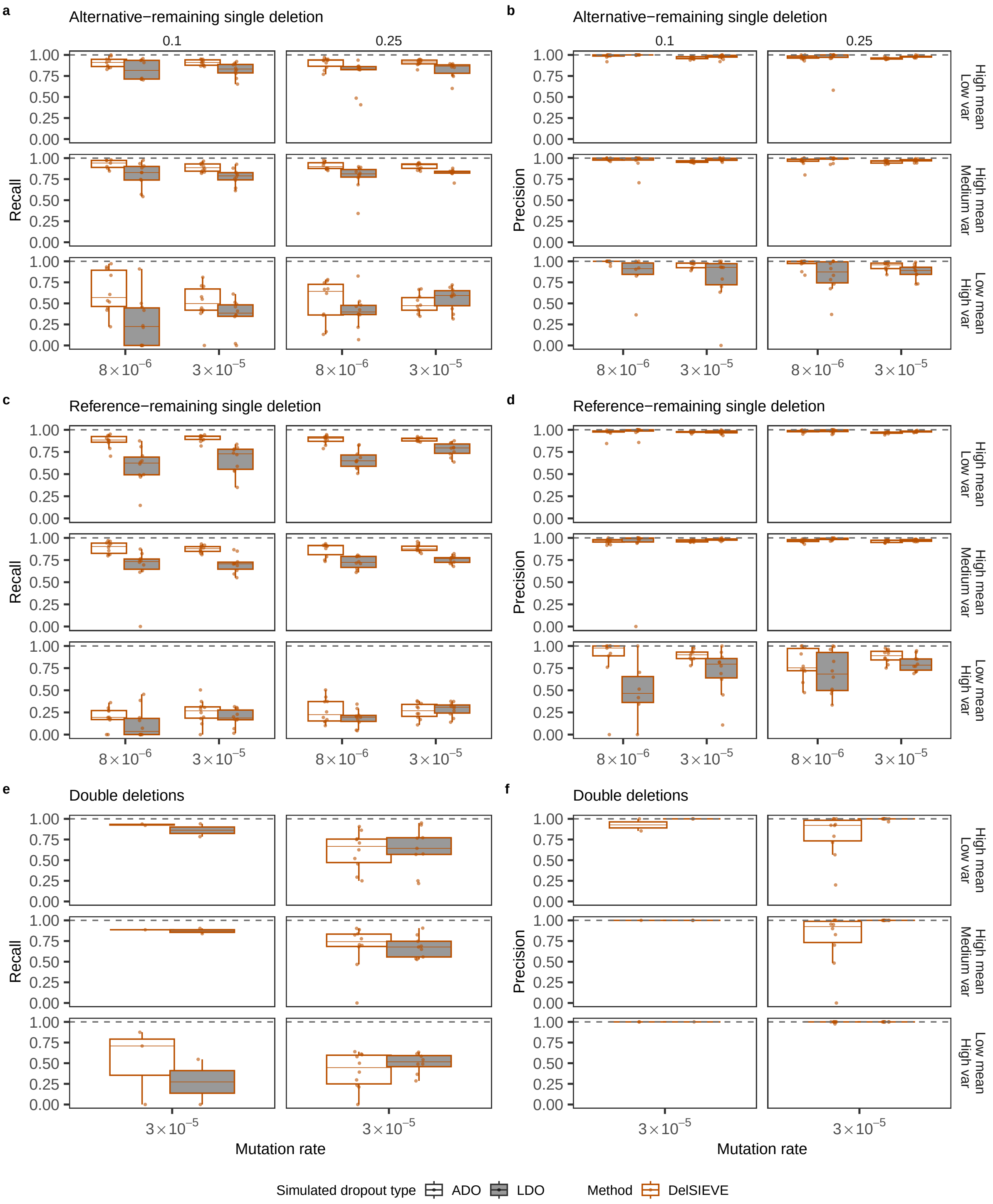
a
Alternative−remaining single deletion
b
Alternative−remaining single deletion
c
Reference−remaining single deletion
d
Reference−remaining single deletion
e
Double deletions
f
Double deletions
0.1
0.25
Recall
Precision
High mean Low var
High mean Medium var
Low mean High var
1.00
0.75
0.50
0.25
0.00
8×10−6
3×10−5
Mutation rate
Simulated dropout type
ADO
LDO
Method
DelSIEVE

**Figure S1** ***(previous page)*****: Recall and precision for the benchmark of calling deletions.** Varying are the mutation rate (the horizontal axis), the relative deletion rate (the vertical strip), the coverage quality (the horizontal strip) and the simulated dropout type (the shaded or blank boxes). Each simulation is repeated $n = 10$ times with each repetition denoted by colored dots. The gray dashed lines represent the optimal values of each metric. Box plots comprise medians, boxes covering the interquartile range (IQR), and whiskers extending to 1.5 times the IQR below and above the box. Data points were removed if the proportion of simulated ground truth was less than 0.1%. Both DelSIEVE and SIEVE were configured to match the dropout mode (ADO or LDO) employed during the simulation process. **a-b**, Box plots of the recall (**a**) and the precision (**b**) for calling alternative-left single deletion. **c-d**, Box plots of the recall (**c**) and the precision (**d**) for calling reference-left single deletion. **e-f**, Box plots of the recall (**e**) and the precision (**f**) for calling double deletions, where the results when mutation rate was $8 \times 10^{-6}$ were omitted as very few double deletions were generated (less than 0.2%).

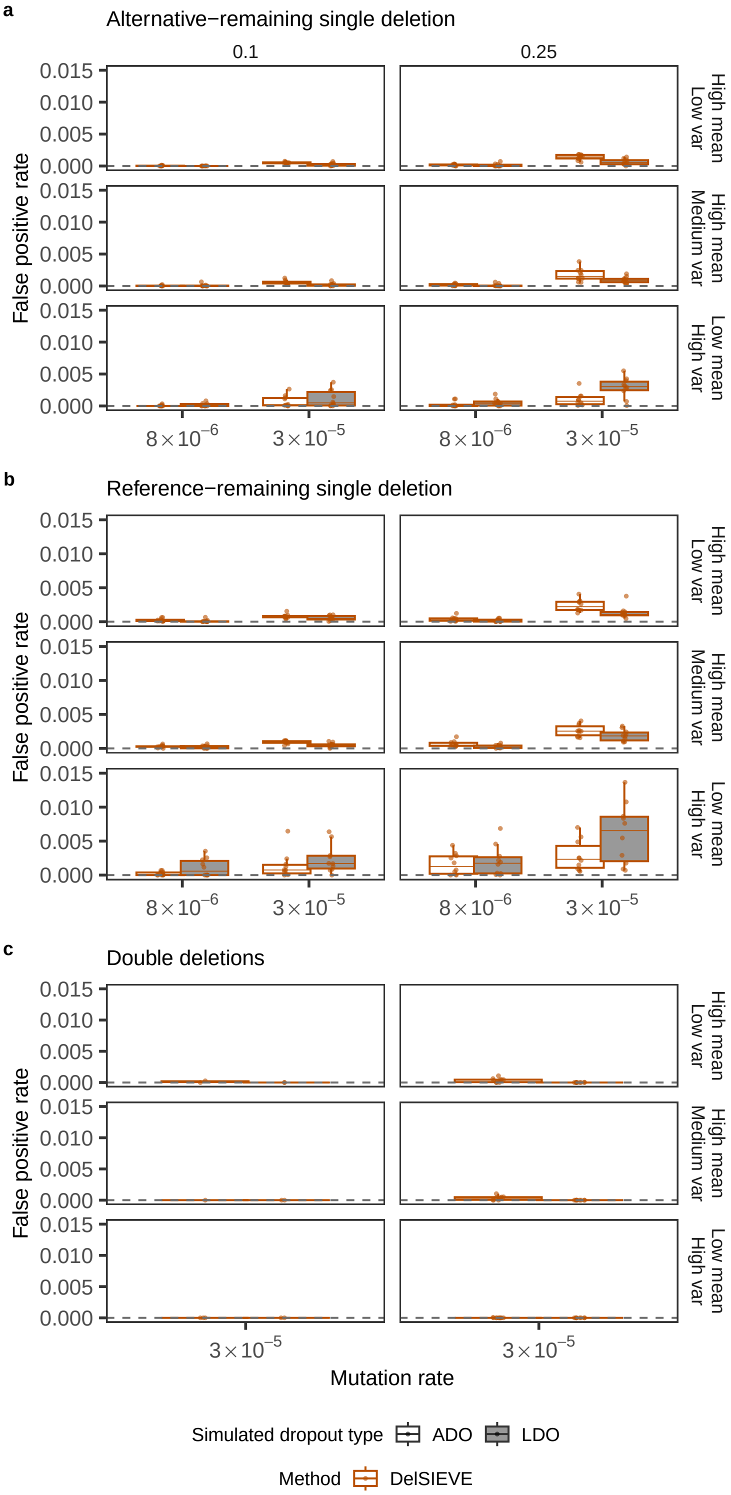
a
Alternative−remaining single deletion
0.1
0.25
False positive rate
0.015
0.010
0.005
0.000
High mean Low var
High mean Medium var
Low mean High var
8×10−6
3×10−5
b
Reference−remaining single deletion
c
Double deletions
Mutation rate
Simulated dropout type
ADO
LDO
Method
DelSIEVE

**Figure S2** ***(previous page)*****: False positive rate (FPR) for the benchmark of calling deletions.** Varying are the mutation rate (the horizontal axis), the relative deletion rate (the vertical strip), the coverage quality (the horizontal strip) and the simulated dropout type (the shaded or blank boxes). Each simulation is repeated $n = 10$ times with each repetition denoted by colored dots. The gray dashed lines represent the optimal values of each metric. Box plots comprise medians, boxes covering the interquartile range (IQR), and whiskers extending to 1.5 times the IQR below and above the box. Data points were removed if the proportion of simulated ground truth was less than 0.1%. Both DelSIEVE and SIEVE were configured to match the dropout mode (ADO or LDO) employed during the simulation process. **a-c**, Box plots of the FPR for calling alternative-left single deletion (**a**), reference-left single deletion (**b**), and double deletions (**c**). The results in **c** when mutation rate was $8 \times 10^{-6}$ were omitted as very few double deletions were generated (less than 0.2%).

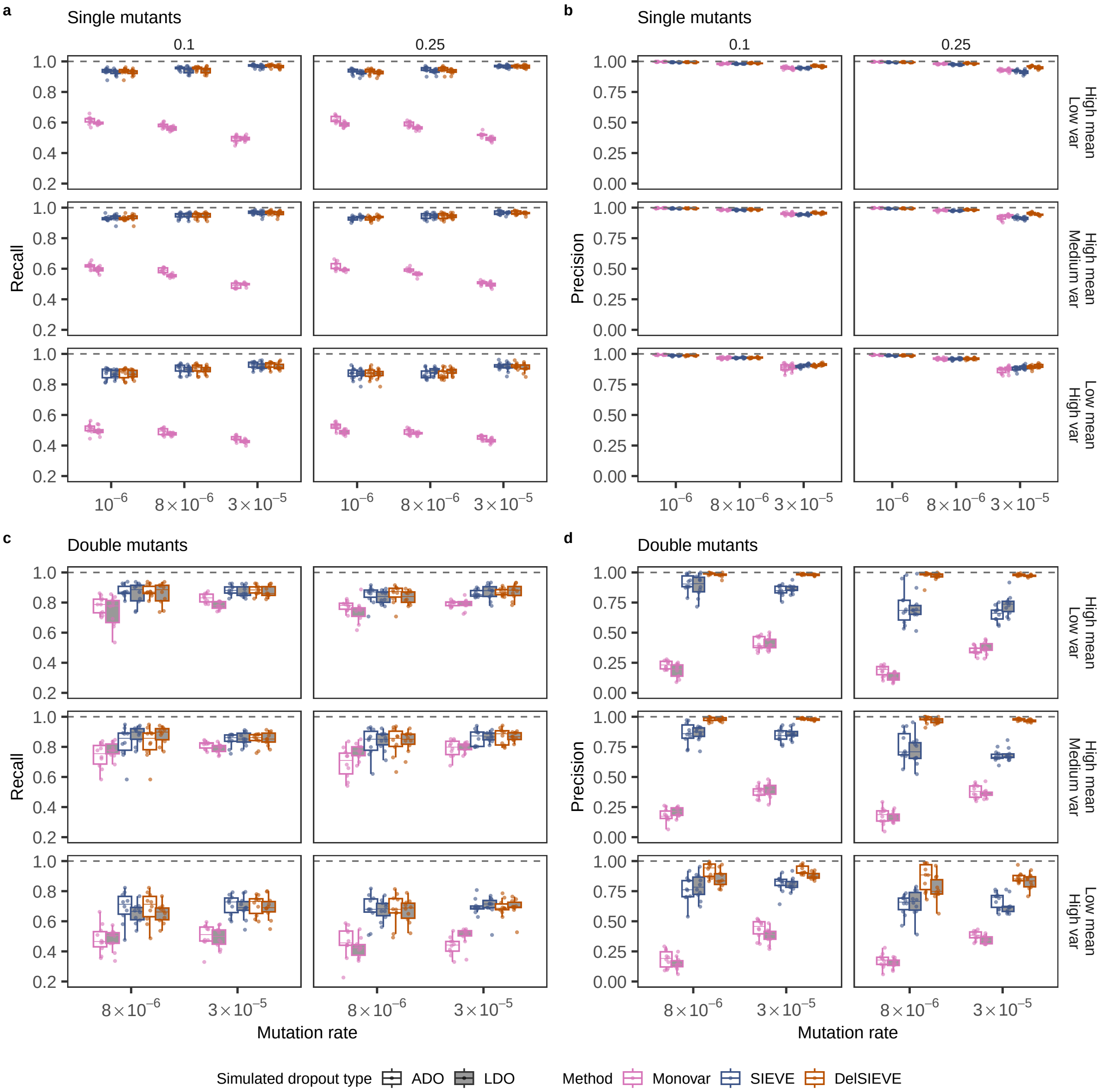


**Figure S3: Recall and precision for the benchmark of calling single and double mutant.** Varying are the mutation rate (the horizontal axis), the relative deletion rate (the vertical strip), the coverage quality (the horizontal strip) and the simulated dropout type (the shaded or blank boxes). Each simulation is repeated $n = 10$ times with each repetition denoted by colored dots. The gray dashed lines represent the optimal values of each metric. Box plots comprise medians, boxes covering the interquartile range (IQR), and whiskers extending to 1.5 times the IQR below and above the box. Both DelSIEVE and SIEVE were configured to match the dropout mode (ADO or LDO) employed during the simulation process. **a-b**, Box plots of the recall (**a**) and the precision (**b**) for calling single mutant. **c-d**, Box plots of the recall (**c**) and the precision (**d**) for calling double mutant, where the results when mutation rate was $10^{-6}$ were omitted as very few double mutant were generated (less than 0.2%).

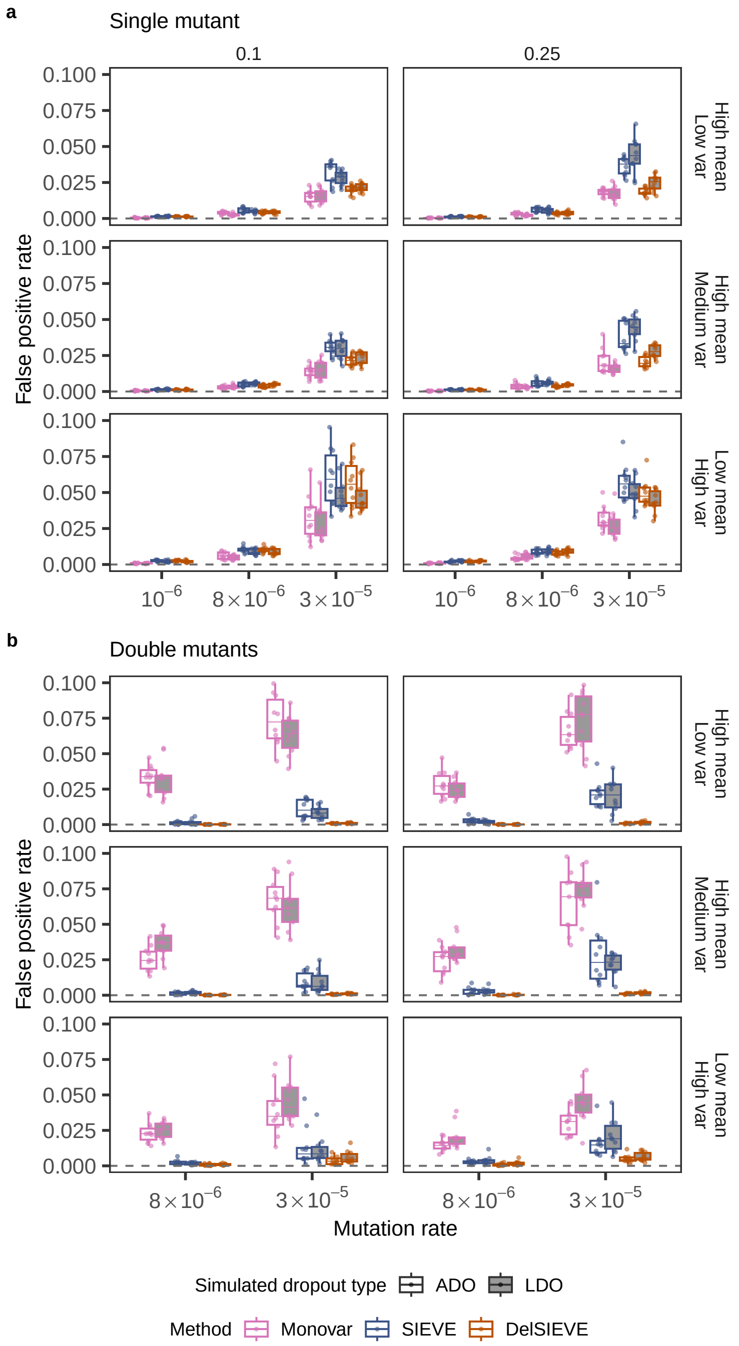


**Figure S4: False positive rate (FPR) for the benchmark of calling single and double mutant.** Varying are the mutation rate (the horizontal axis), the relative deletion rate (the vertical strip), the coverage quality (the horizontal strip) and the simulated dropout type (the shaded or blank boxes). Each simulation is repeated $n = 10$ times with each repetition denoted by colored dots. The gray dashed lines represent the optimal values of each metric. Box plots comprise medians, boxes covering the interquartile range (IQR), and whiskers extending to 1.5 times the IQR below and above the box. Both DelSIEVE and SIEVE were configured to match the dropout mode (ADO or LDO) employed during the simulation process. **a-b**, Box plots of the FPR for calling single mutant (**a**) and double mutant (**b**). The results in **b** when mutation rate was $10^{-6}$ were omitted as very few double mutant were generated (less than 0.2%).

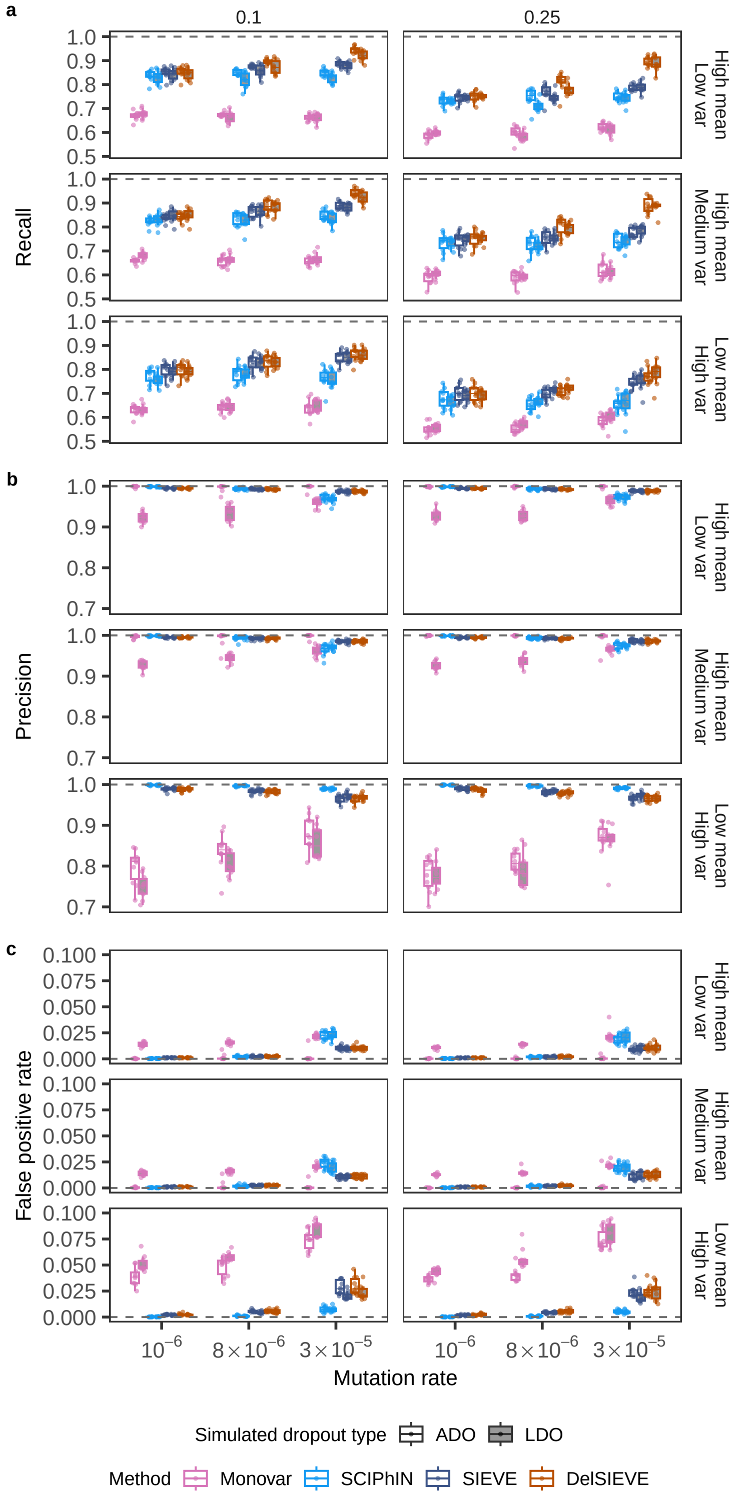


**Figure S5: Recall, precision and false positive rate (FPR) for the benchmark of the DelSIEVE model.** Varying are the mutation rate (the horizontal axis), the relative deletion rate (the vertical strip), the coverage quality (the horizontal strip), and the simulated dropout type (the shaded or blank boxes). Each simulation is repeated $n = 10$ times, with each repetition denoted by colored dots. The gray dashed lines represent the optimal values of each metric. Box plots comprise medians, boxes covering the interquartile range (IQR), and whiskers extending to 1.5 times the IQR below and above the box. Data points were removed if the proportion of simulated ground truth was less than 0.1%. Both DelSIEVE and SIEVE were configured to match the dropout mode (ADO or LDO) employed during the simulation process. **a-c**, Box plots of the recall (**a**), precision (**b**) and FPR (**c**) for calling general "mutations", composed of all genotypes other than wildtype.

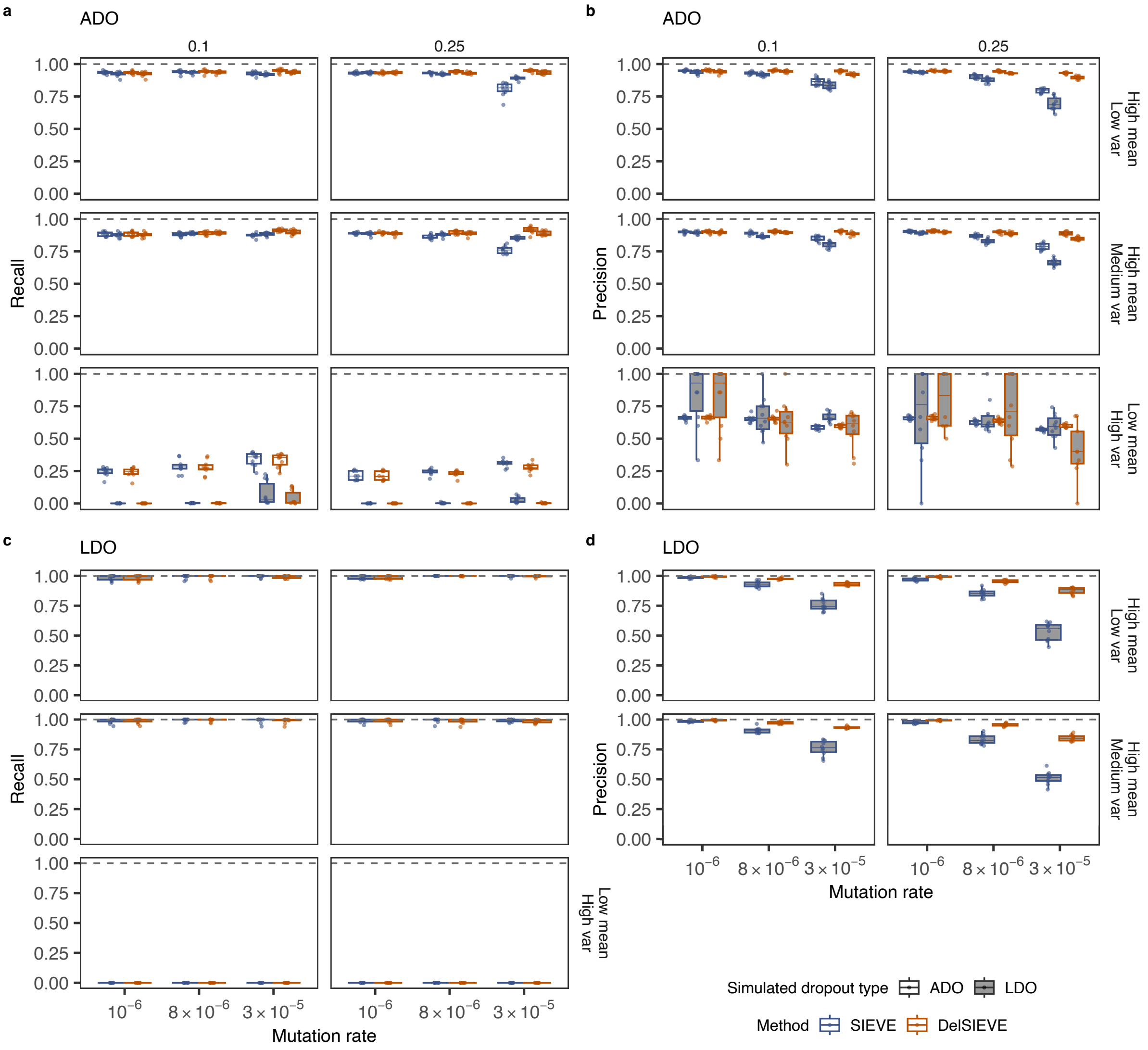


**Figure S6: Recall and precision for the benchmark of calling ADO and LDO.** Varying are the mutation rate (the horizontal axis), the relative deletion rate (the vertical strip), the coverage quality (the horizontal strip) and the simulated dropout type (the shaded or blank boxes). Each simulation is repeated $n = 10$ times with each repetition denoted by colored dots. The gray dashed lines represent the optimal values of each metric. Box plots comprise medians, boxes covering the interquartile range (IQR), and whiskers extending to 1.5 times the IQR below and above the box. Both DelSIEVE and SIEVE were configured to match the dropout mode (ADO or LDO) employed during the simulation process. **a-b**, Box plots of the recall (**a**) and the precision (**b**) for calling single ADO. **c-d**, Box plots of the recall (**c**) and the precision (**d**) for calling locus dropout, where the precision were unavailable in **d** when data was of low coverage quality due to zero called locus dropout.

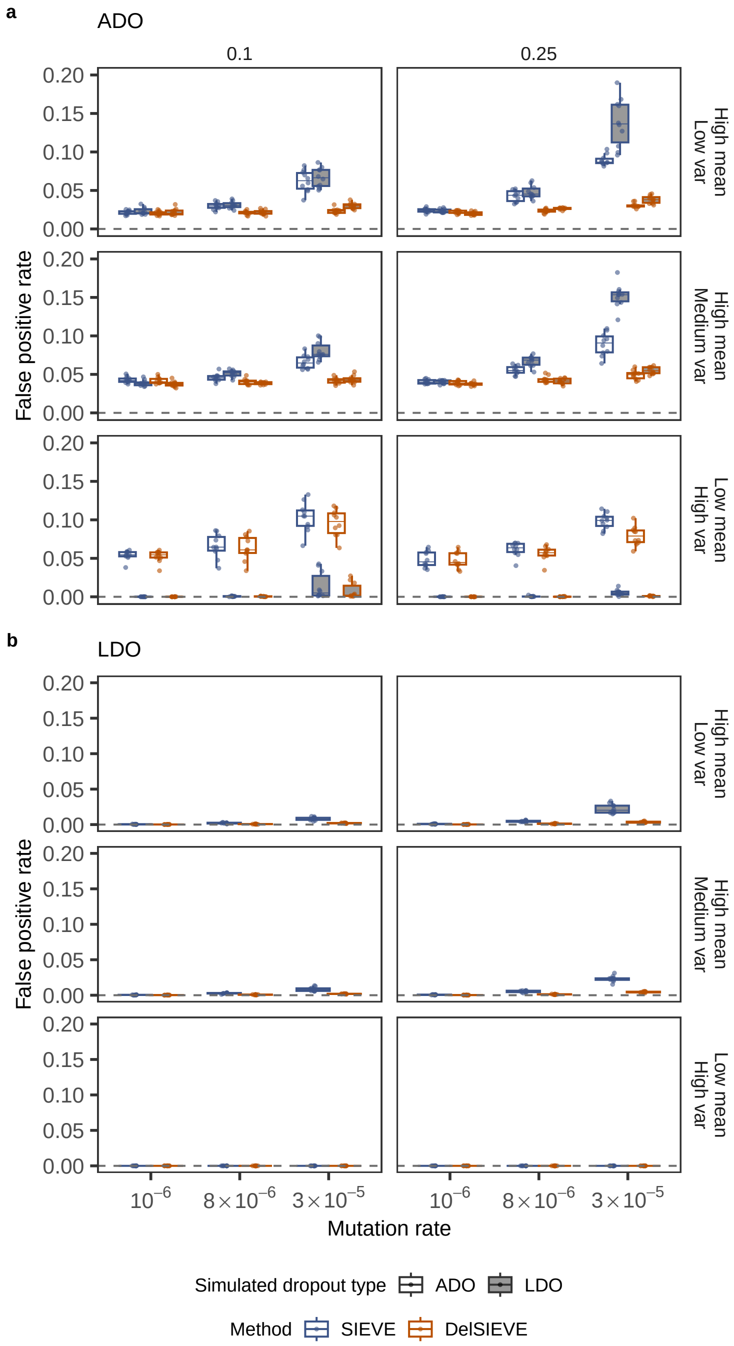


**Figure S7: False positive rate (FPR) for the benchmark of calling ADO and LDO.** Varying are the mutation rate (the horizontal axis), the relative deletion rate (the vertical strip), the coverage quality (the horizontal strip) and the simulated dropout type (the shaded or blank boxes). Each simulation is repeated $n = 10$ times with each repetition denoted by colored dots. The gray dashed lines represent the optimal values of each metric. Box plots comprise medians, boxes covering the interquartile range (IQR), and whiskers extending to 1.5 times the IQR below and above the box. Both DelSIEVE and SIEVE were configured to match the dropout mode (ADO or LDO) employed during the simulation process. **a-b**, Box plots of the FPR for calling single ADO (**a**) and locus dropout (**b**).

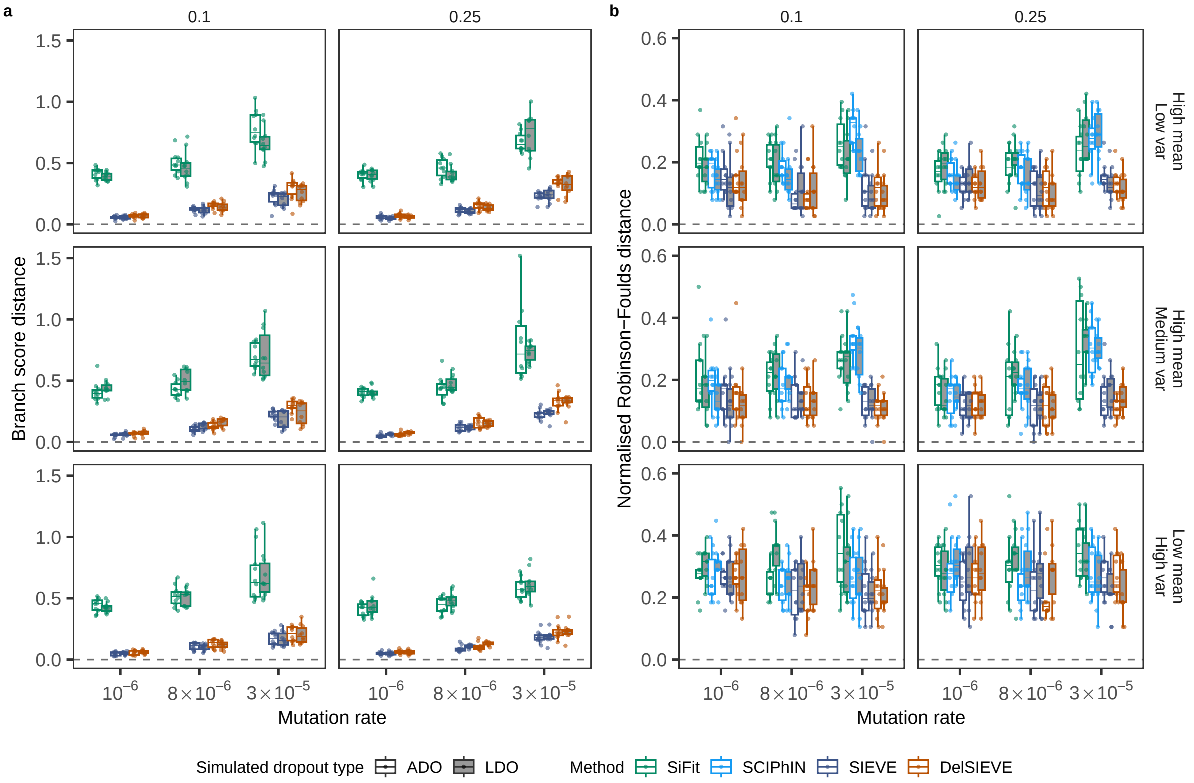


**Figure S8: Benchmark of tree inference accuracy.** Varying are the mutation rate (the horizontal axis), the relative deletion rate (the vertical strip), the coverage quality (the horizontal strip) and the simulated dropout type (the shaded or blank boxes). Each simulation is repeated $n = 10$ times with each repetition denoted by colored dots. The gray dashed lines represent the optimal values of each metric. Box plots comprise medians, boxes covering the interquartile range (IQR), and whiskers extending to 1.5 times the IQR below and above the box. Both DelSIEVE and SIEVE were configured to match the dropout mode (ADO or LDO) employed during the simulation process. **a-b**, Box plots of the BS distance where the branch lengths are taken into account (**a**) and the normalized RF distance where only tree topology is considered (**b**).

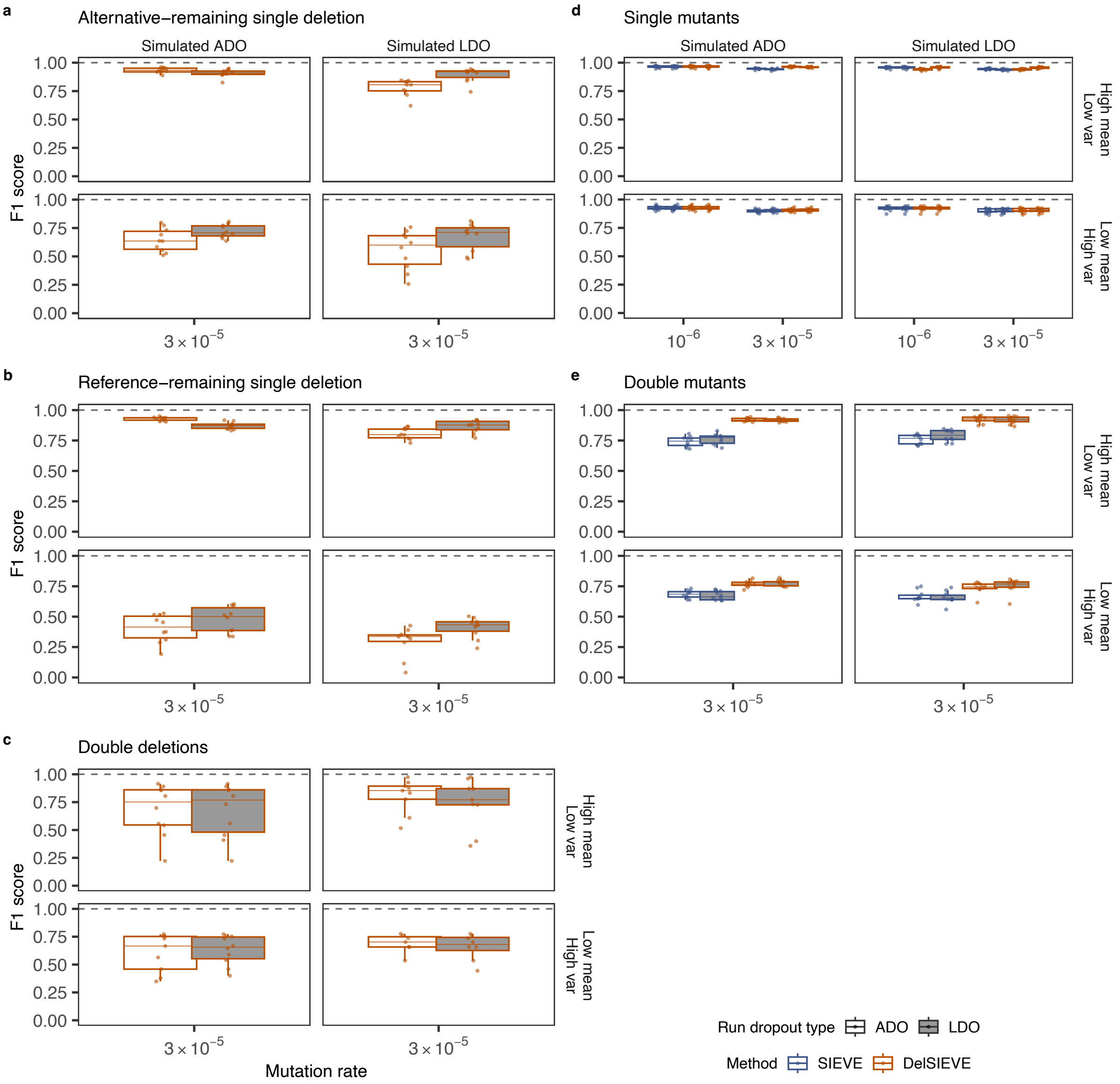


**Figure S9: F1 score for the benchmark of calling all genotypes other than wildtype.** Varying are the mutation rate (the horizontal axis), the dropout type used to simulate the data (the vertical strip), the coverage quality (the horizontal strip), and the dropout type used to configure DelSIEVE and SIEVE (the shaded or blank boxes). Each simulation is repeated $n = 10$ times, with each repetition denoted by colored dots. The gray dashed lines represent the optimal values of each metric. Box plots comprise medians, boxes covering the interquartile range (IQR), and whiskers extending to 1.5 times the IQR below and above the box. Both DelSIEVE and SIEVE were run under ADO and LDO modes, regardless of that used to simulate the data. Data points were removed if the proportion of simulated ground truth was less than 0.1%. **a-e**, Box plots of the F1 score for calling alternative-left single deletion (**a**), reference-left single deletion (**b**), double deletions (**c**), single mutant (**d**), and double mutant (**e**).

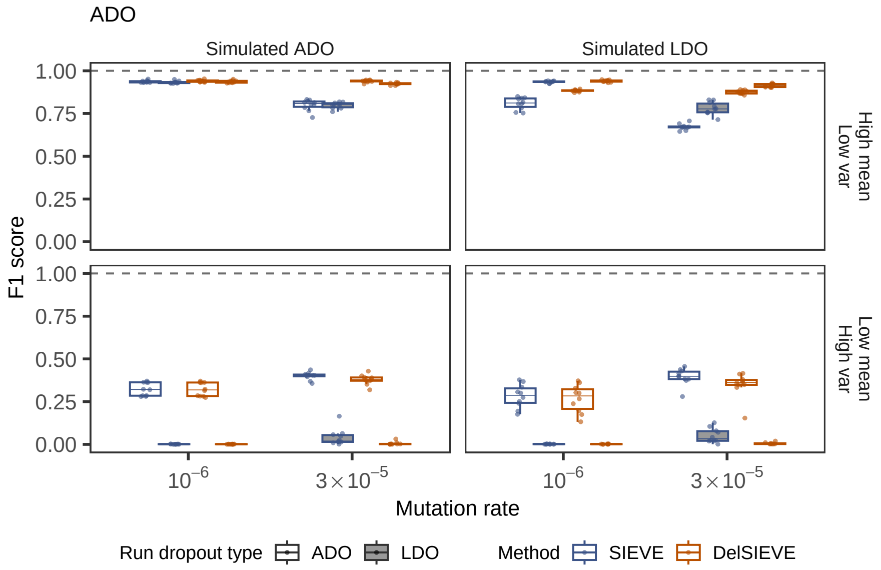


**Figure S10: F1 score for the benchmark of calling ADO.** Varying are the mutation rate (the horizontal axis), the dropout type used to simulate the data (the vertical strip), the coverage quality (the horizontal strip), and the dropout type used to configure DelSIEVE and SIEVE (the shaded or blank boxes). Each simulation is repeated $n = 10$ times, with each repetition denoted by colored dots. The gray dashed lines represent the optimal values of each metric. Box plots comprise medians, boxes covering the interquartile range (IQR), and whiskers extending to 1.5 times the IQR below and above the box. Both DelSIEVE and SIEVE were run under ADO and LDO mode, regardless of that used to simulate the data.

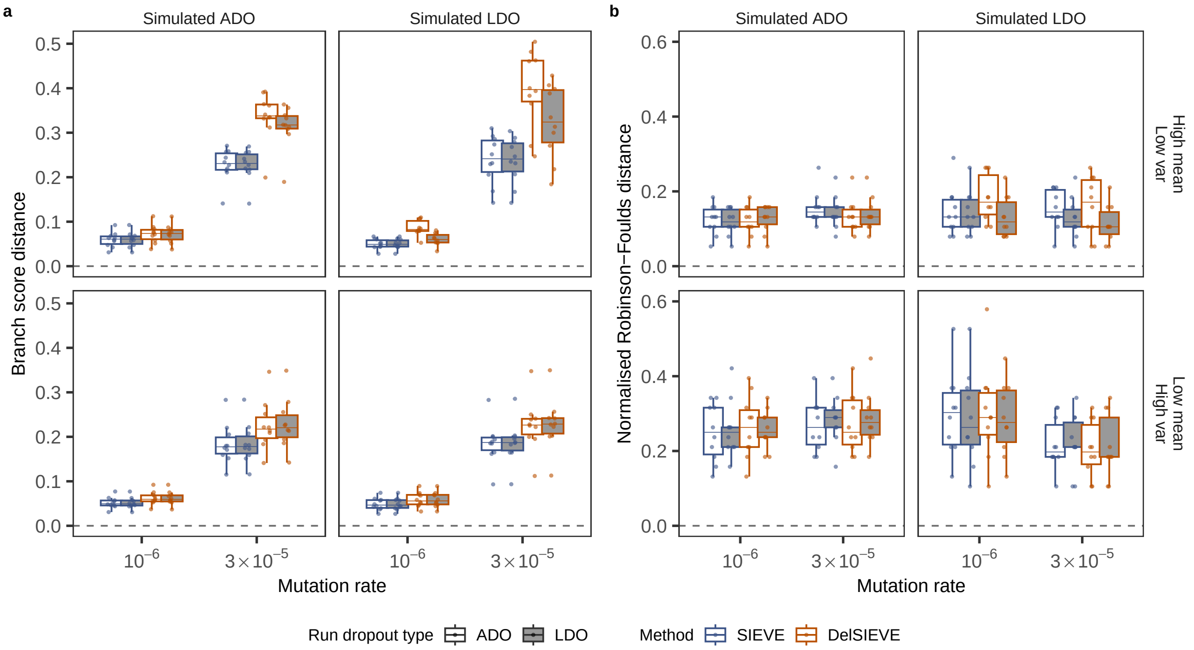


**Figure S11: Benchmark of tree inference accuracy.** Varying are the mutation rate (the horizontal axis), the dropout type used to simulate the data (the vertical strip), the coverage quality (the horizontal strip), and the dropout type used to configure DelSIEVE and SIEVE (the shaded or blank boxes). Each simulation is repeated $n = 10$ times, with each repetition denoted by colored dots. The gray dashed lines represent the optimal values of each metric. Box plots comprise medians, boxes covering the interquartile range (IQR), and whiskers extending to 1.5 times the IQR below and above the box. Both DelSIEVE and SIEVE were run under ADO and LDO mode, regardless of that used to simulate the data. **a-b**, Box plots of the BS distance where the branch lengths are taken into account (**a**) and the normalized RF distance where only tree topology is considered (**b**).

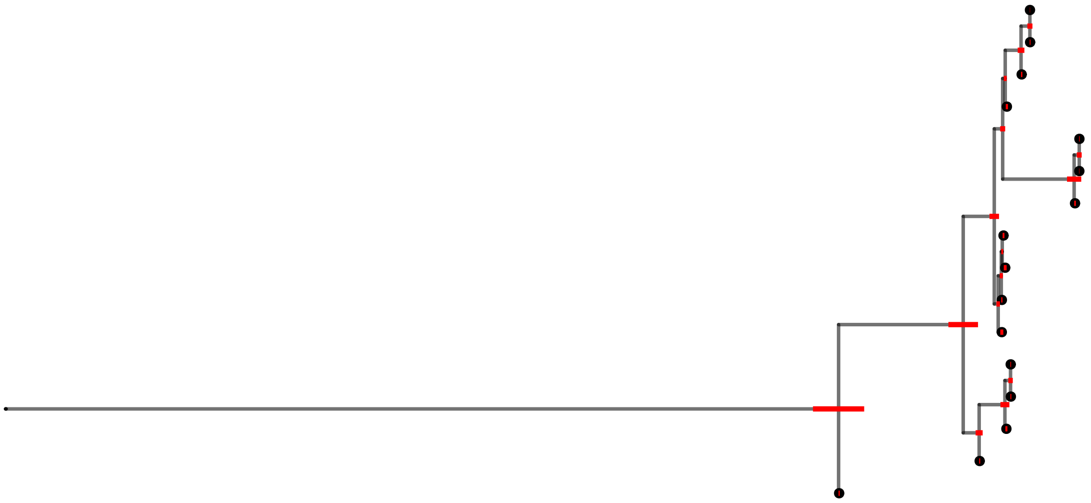

**Figure S12: Illustration of branch lengths of the phylogenetic tree inferred from TNBC16 [38] by DelSIEVE.** Shown is exactly the same tree as in Figure 4, except that cell names, subclone posterior probabilities and gene annotations are removed and no branches are folded. Red bars annotated to internal nodes except the root are the 95% HPD intervals of the corresponding branch lengths.

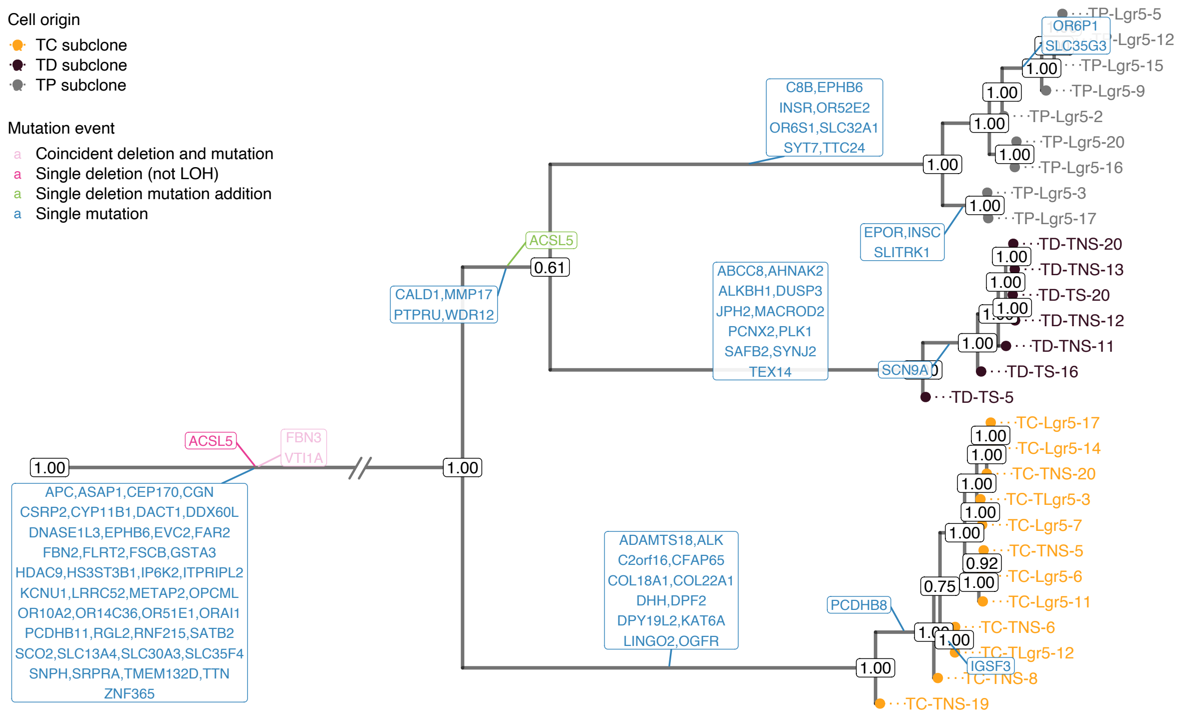


**Figure S13: Results of phylogenetic inference for the CRC28 [32] dataset.** Shown is DelSIEVE's maximum clade credibility tree. Tumor cell names are annotated to the leaves of the tree. The exceptionally long trunk has been folded (marked by slashes). Cells are colored according to the corresponding biopsies. The numbers at each node represent the posterior probabilities (threshold $p > 0.5$). At each branch, depicted in different colors are non-synonymous genes that are either CRC-related single mutations (in blue) or other mutation events (in other colors).

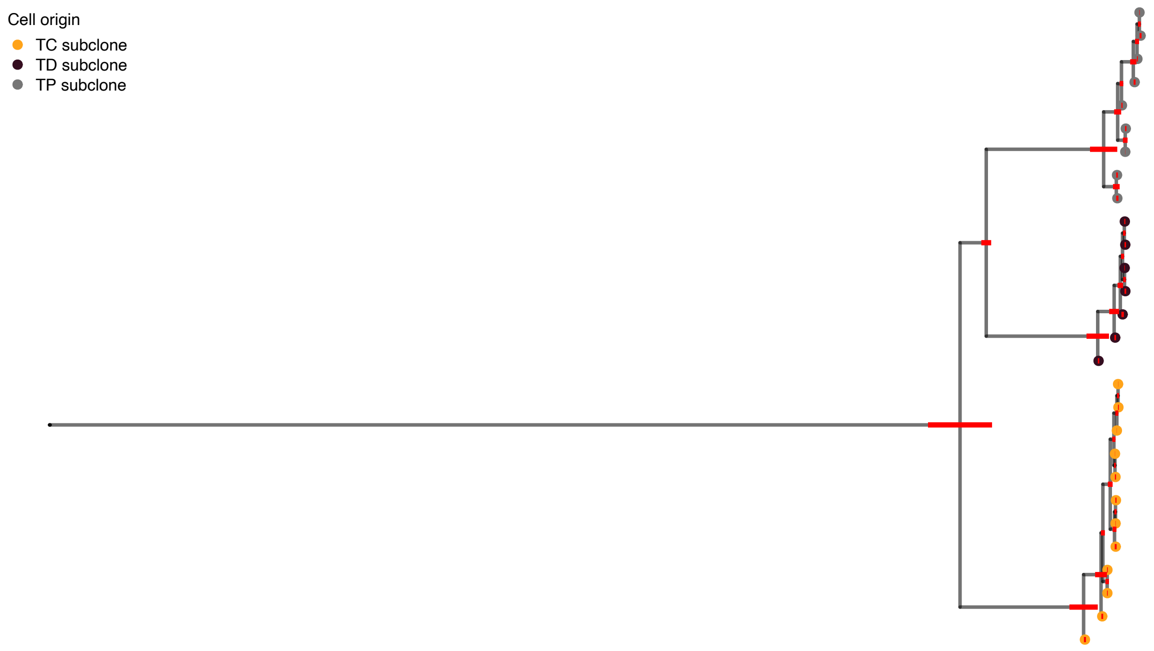


**Figure S14: Illustration of branch lengths of the phylogenetic tree inferred from CRC28 [32] by DelSIEVE.** Shown is exactly the same tree as in Figure S13, except that cell names, subclone posterior probabilities and gene annotations are removed and no branches are folded. Red bars annotated to internal nodes except the root are the 95% HPD intervals of the corresponding branch lengths.

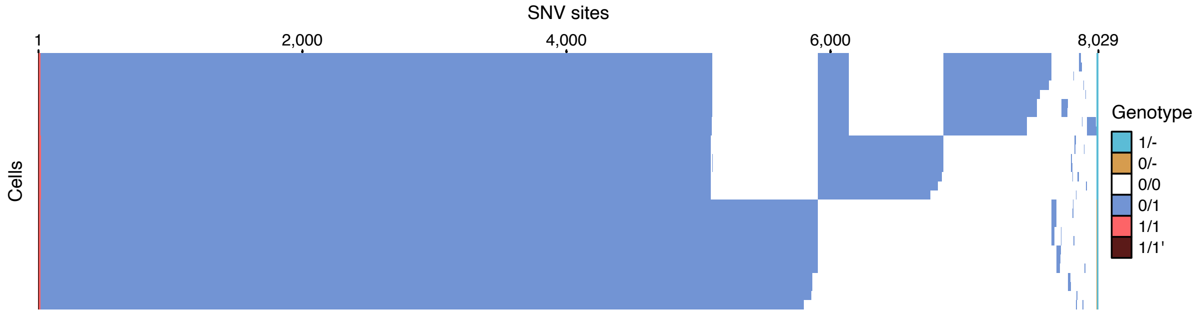


**Figure S15: Results of variant calling for the CRC28 dataset [32].** Cells in the row are in the same order as that of leaves in the phylogenetic tree in Figure S13.

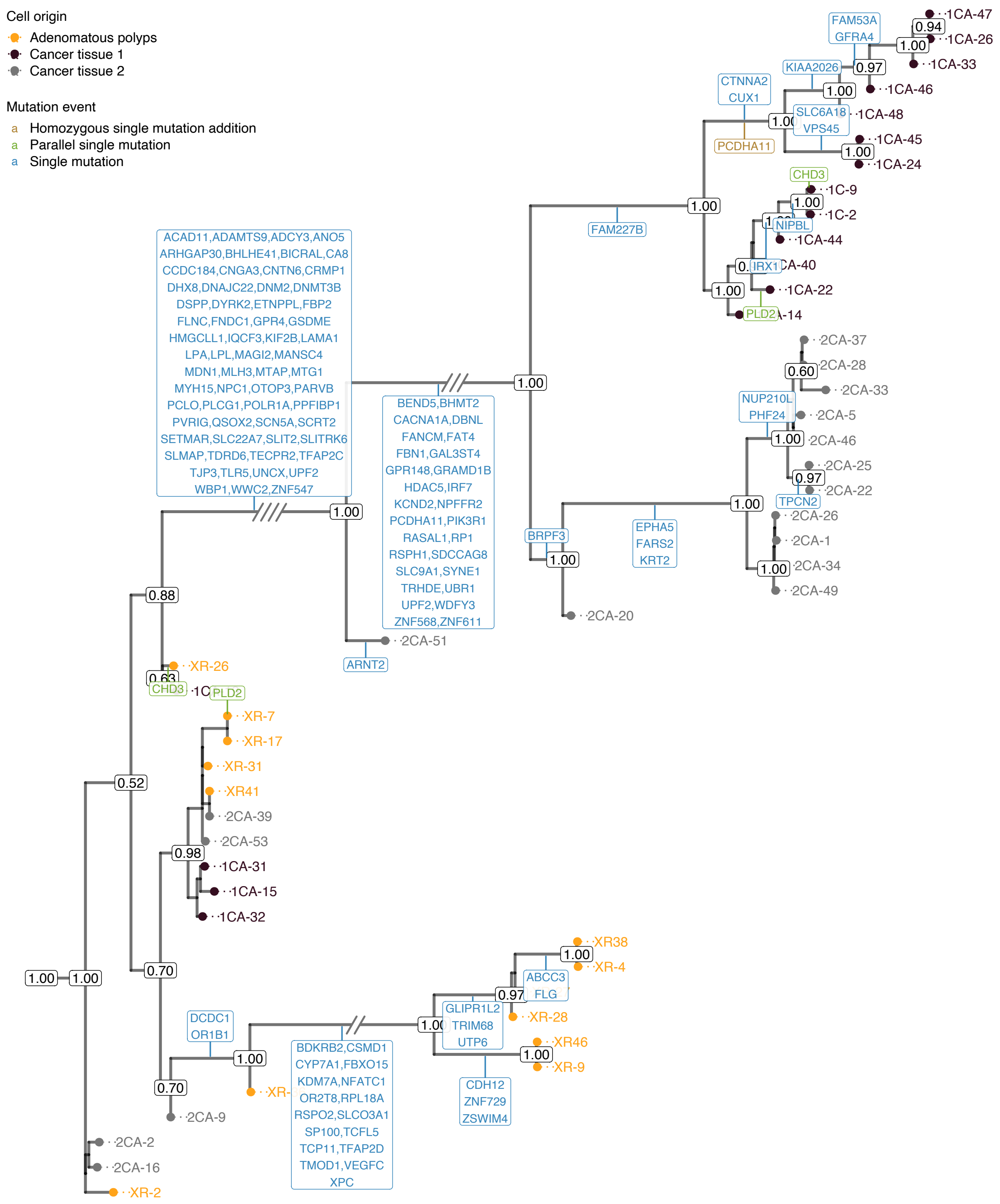


**Figure S16: Results of phylogenetic inference for the CRC48 dataset [42].** Shown is DelSIEVE's maximum clade credibility tree. Tumor cell names are annotated to the leaves of the tree. Three exceptionally long branches are folded with the number of slashes proportional to the branch lengths. Cells are colored according to the corresponding biopsies. The numbers at each node represent the posterior probabilities (threshold $p > 0.5$). At each branch, depicted in different colors are non-synonymous genes that are either CRC-related single mutations (in blue) or other mutation events (in other colors).

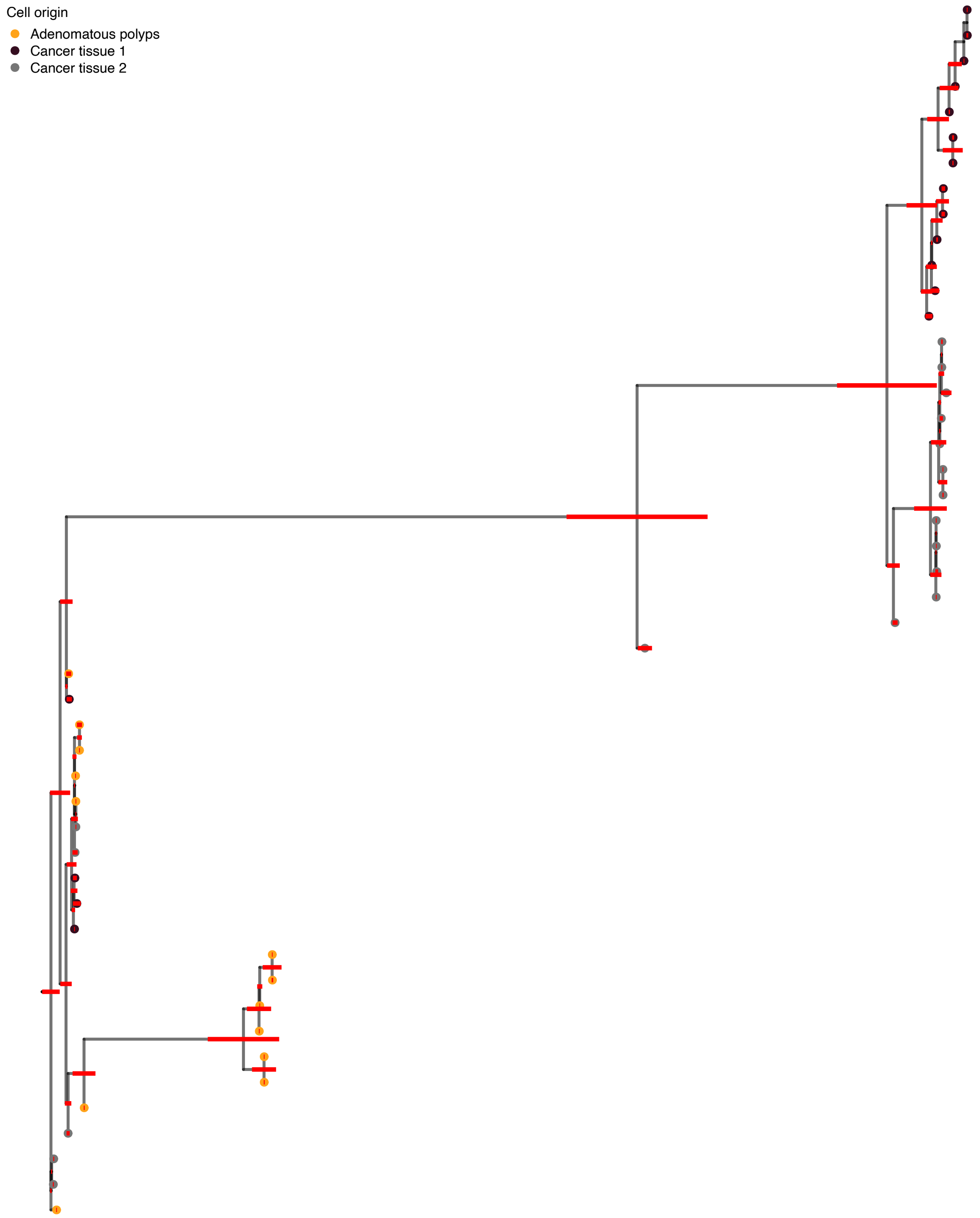


**Figure S17: Illustration of branch lengths of the phylogenetic tree inferred from CRC48 [42] by DelSIEVE.** Shown is exactly the same tree as in Figure S16, except that cell names, subclone posterior probabilities and gene annotations are removed and no branches are folded. Red bars annotated to internal nodes except the root are the 95% HPD intervals of the corresponding branch lengths.

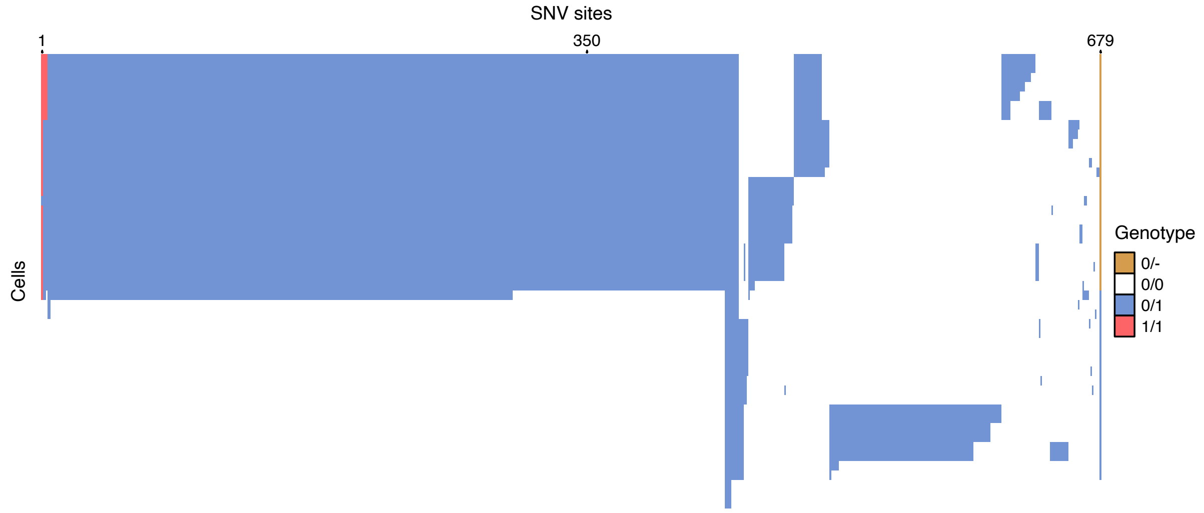


**Figure S18: Results of variant calling for the CRC48 dataset [42].** Cells in the row are in the same order as that of leaves in the phylogenetic tree in Figure S16.

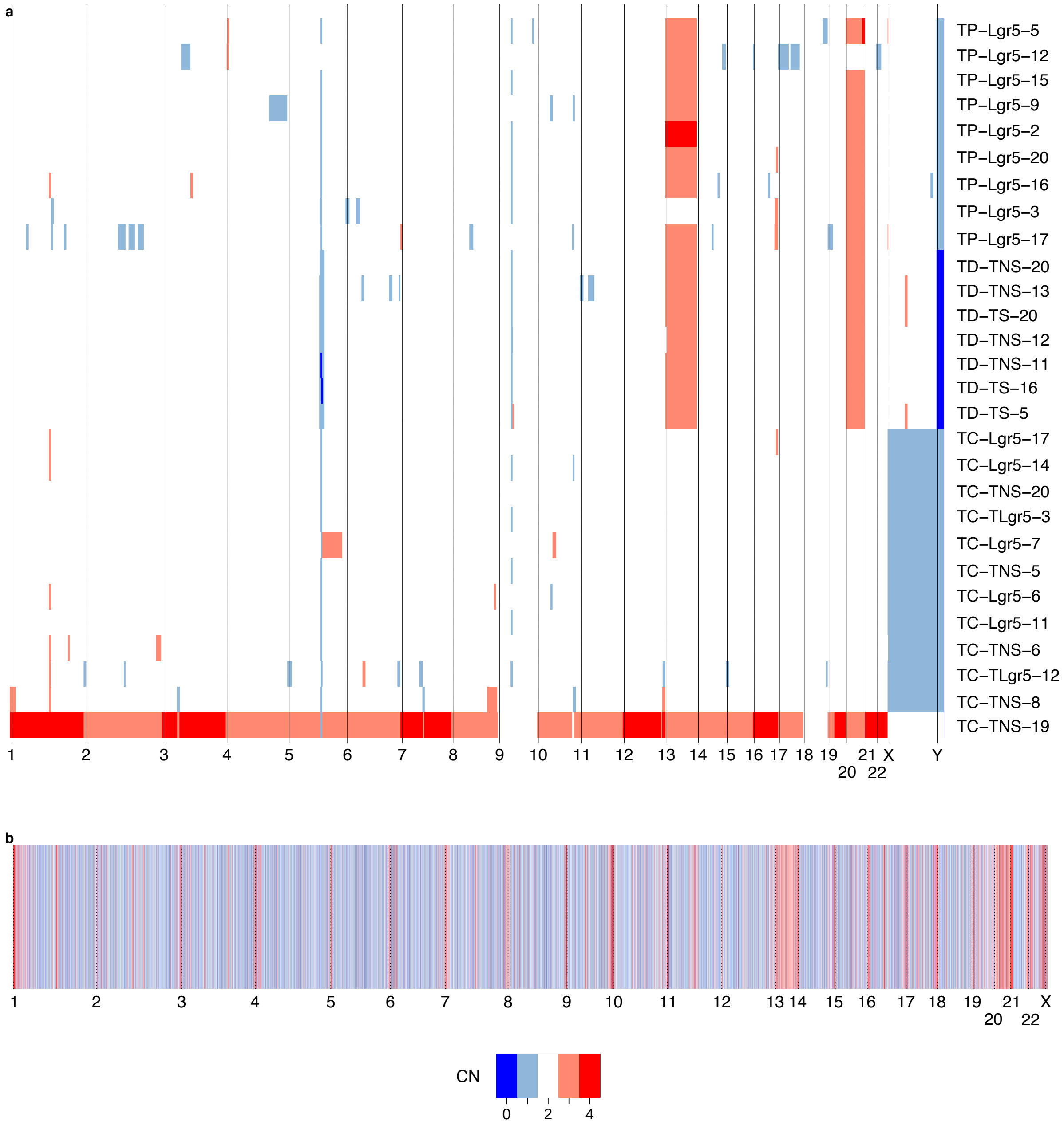


**Figure S19: Heatmaps of copy numbers (CNs) across whole genome of cells in CRC28 [32] reported by Ginkgo and Sequenza.** The horizontal axis represents the indices of chromosomesm, and the vertical axis represents either single cells (in **a**, by Ginkgo) or bulk-seq sample (in **b**, by Sequenza).

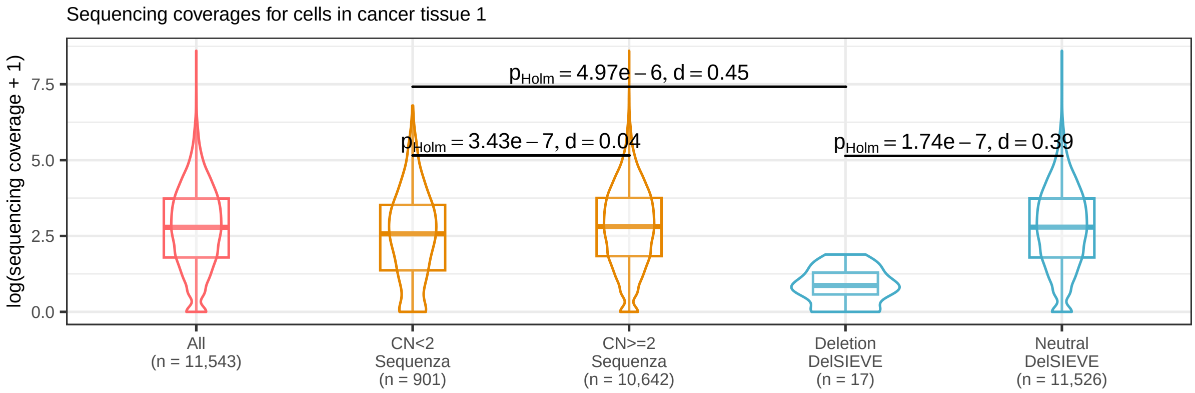


**Figure S20: Results of subclone-wise sequencing coverage comparison for cells in cancer tissue 1 of CRC48 [42] between DelSIEVE and Sequenza.** Compared were the sites shared between the input data of both methods. The resolution of variant calling was subclone-wise in order to conduct a fair comparison. For Sequenza, sites were divided into two groups with copy number (CN) $< 2$ and $\geq 2$, respectively. For DelSIEVE, sites were divided into two groups, one with deletions, the other copy neutral. Sequencing coverage transformed with $\log p1$ across those cells in the subclone at all sites were plotted for reference. In each group, the violin and the box plots showed the distribution of the sequencing coverage. The total number of dots in each group, which was the product of the number of cells (17) and the number of sites in each group, was marked with $n$ on the horizontal axis. Box plots comprise medians, boxes covering the interquartile range (IQR), and whiskers extending to 1.5 times the IQR below and above the box. Within- and between-group comparisons were conducted between CN $< 2$ and $\geq 2$ of Sequenza, between deletions and copy neutral of DelSIEVE, and between CN $< 2$ of Sequenza and deletions of DelSIEVE. Each comparison was conducted on the sequencing coverage on the original scale, showing the result of Mann-Whitney U test, with the p-value corrected by Holm–Bonferroni method and the absolute value of the effect size (Cohen's d).

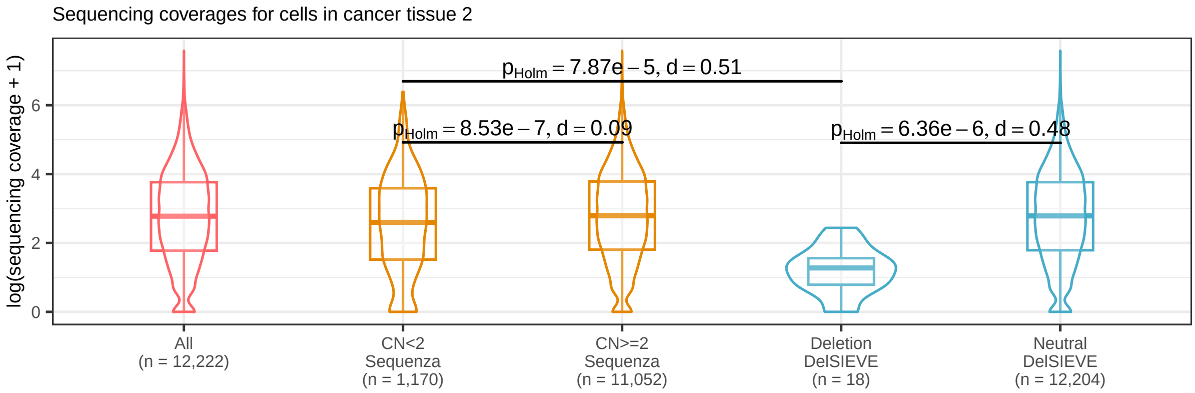


**Figure S21: Results of subclone-wise sequencing coverage comparison for cells in cancer tissue 2 of CRC48 [42] between DelSIEVE and Sequenza.** Compared were the sites shared between the input data of both methods. The resolution of variant calling was subclone-wise in order to conduct a fair comparison. For Sequenza, sites were divided into two groups with copy number (CN) $< 2$ and $\geq 2$, respectively. For DelSIEVE, sites were also divided into two groups, one with deletions, the other copy neutral. Sequencing coverage transformed with $\log p1$ across those cells in the subclone at all sites were plotted for reference. In each group, the violin and the box plots showed matched the color of the method and the distribution of the sequencing coverage. The total number of dots in each group, which was the product of the number of cells (18) and the number of sites in each group, was marked with $n$ on the horizontal axis. Box plots comprise medians, boxes covering the interquartile range (IQR), and whiskers extending to 1.5 times the IQR below and above the box. Within- and between-group comparisons were conducted between CN $< 2$ and $\geq 2$ of Sequenza, between deletions and copy neutral of DelSIEVE, and between CN $< 2$ of Sequenza and deletions of DelSIEVE. Each comparison was conducted on the sequencing coverage on the original scale, showing the result of Mann-Whitney U test, with the p-value corrected by Holm–Bonferroni method and the absolute value of the effect size (Cohen's d).

### Supplementary tables

**Table S1: Definition of the distribution of the number of sequenced alleles ($\alpha_{ij}$) conditional on the true genotype ($g_{ij}$) and on the allelic ADO rate ($\theta_A$) under the ADO mode for DelSIEVE.**

| $\alpha_{ij}$ | $g_{ij}$ | ADO occurred | $P(\alpha_{ij} \mid g_{ij}, \theta_A)$ |
|---|---|---|---|
| 1 | 0/0 | Yes | $\theta_A$ |
| 2 | 0/0 | No | $1 - \theta_A$ |
| 1 | 0/1 | Yes | $\theta_A$ |
| 2 | 0/1 | No | $1 - \theta_A$ |
| 1 | 1/1 | Yes | $\theta_A$ |
| 2 | 1/1 | No | $1 - \theta_A$ |
| 1 | $1/1'$ | Yes | $\theta_A$ |
| 2 | $1/1'$ | No | $1 - \theta_A$ |
| 0 | 0/- | Yes | $\theta_A/2$ |
| 1 | 0/- | No | $1 - \theta_A/2$ |
| 0 | 1/- | Yes | $\theta_A/2$ |
| 1 | 1/- | No | $1 - \theta_A/2$ |
| 0 | - | No | 1 |
| | Others | | 0 |

**Table S2: Definition of the distribution of $\alpha_{ij}$ conditional on $g_{ij}$ and $\theta_L$ under LDO mode for DelSIEVE.** Definition of the distribution of the number of sequenced alleles ($\alpha_{ij}$) conditional on the true genotype ($g_{ij}$) and on the allelic ADO rate ($\theta_L$) under the LDO mode for DelSIEVE.

| $\alpha_{ij}$ | $g_{ij}$ | Number of alleles dropped out | $P(\alpha_{ij} \mid g_{ij}, \theta_L)$ |
|---|---|---|---|
| 0 | 0/0 | 2 | $\theta_L^2$ |
| 1 | 0/0 | 1 | $2\theta_L(1-\theta_L)$ |
| 2 | 0/0 | 0 | $(1-\theta_L)^2$ |
| 0 | 0/1 | 2 | $\theta_L^2$ |
| 1 | 0/1 | 1 | $2\theta_L(1-\theta_L)$ |
| 2 | 0/1 | 0 | $(1-\theta_L)^2$ |
| 0 | 1/1 | 2 | $\theta_L^2$ |
| 1 | 1/1 | 1 | $2\theta_L(1-\theta_L)$ |
| 2 | 1/1 | 0 | $(1-\theta_L)^2$ |
| 0 | 1/1′ | 2 | $\theta_L^2$ |
| 1 | 1/1′ | 1 | $2\theta_L(1-\theta_L)$ |
| 2 | 1/1′ | 0 | $(1-\theta_L)^2$ |
| 0 | 0/- | 1 | $\theta_L$ |
| 1 | 0/- | 0 | $1-\theta_L$ |
| 0 | 1/- | 1 | $\theta_L$ |
| 1 | 1/- | 0 | $1-\theta_L$ |
| 0 | - | 0 | 1 |
| | Others | | 0 |

**Table S3: Evolutionary rate matrix used in the simulator to generate the simulated data for DelSIEVE.** Genotypes are encoded with nucleotides rather than numbers. $d$ is the deletion rate measured relatively to the mutation rate. The diagonal elements are denoted by dots, and have negative values equal to the sum of the other entries in the same row, ensuring that the sum of each row equals zero.

| | A/A | A/C | A/G | A/T | C/C | C/G | C/T | G/G | G/T | T/T | A/- | C/- | G/- | T/- | - |
|---|---|---|---|---|---|---|---|---|---|---|---|---|---|---|---|
| A/A | . | $1/3$ | $1/3$ | $1/3$ | 0 | 0 | 0 | 0 | 0 | 0 | $d$ | 0 | 0 | 0 | 0 |
| A/C | $1/6$ | . | $1/6$ | $1/6$ | $1/6$ | $1/6$ | $1/6$ | 0 | 0 | 0 | $d/2$ | $d/2$ | 0 | 0 | 0 |
| A/G | $1/6$ | $1/6$ | . | $1/6$ | 0 | $1/6$ | 0 | $1/6$ | $1/6$ | 0 | $d/2$ | 0 | $d/2$ | 0 | 0 |
| A/T | $1/6$ | $1/6$ | $1/6$ | . | 0 | 0 | $1/6$ | 0 | $1/6$ | $1/6$ | $d/2$ | 0 | 0 | $d/2$ | 0 |
| C/C | 0 | $1/3$ | 0 | 0 | . | $1/3$ | $1/3$ | 0 | 0 | 0 | 0 | $d$ | 0 | 0 | 0 |
| C/G | 0 | $1/6$ | $1/6$ | 0 | $1/6$ | . | $1/6$ | $1/6$ | $1/6$ | 0 | 0 | $d/2$ | $d/2$ | 0 | 0 |
| C/T | 0 | $1/6$ | 0 | $1/6$ | $1/6$ | $1/6$ | . | 0 | $1/6$ | $1/6$ | 0 | $d/2$ | 0 | $d/2$ | 0 |
| G/G | 0 | 0 | $1/3$ | 0 | 0 | $1/3$ | 0 | . | $1/3$ | 0 | 0 | 0 | $d$ | 0 | 0 |
| G/T | 0 | 0 | $1/6$ | $1/6$ | 0 | $1/6$ | $1/6$ | $1/6$ | . | $1/6$ | 0 | 0 | $d/2$ | $d/2$ | 0 |
| T/T | 0 | 0 | 0 | $1/3$ | 0 | 0 | $1/3$ | 0 | $1/3$ | . | 0 | 0 | 0 | $d$ | 0 |
| A/- | 0 | 0 | 0 | 0 | 0 | 0 | 0 | 0 | 0 | 0 | . | $1/6$ | $1/6$ | $1/6$ | $d/2$ |
| C/- | 0 | 0 | 0 | 0 | 0 | 0 | 0 | 0 | 0 | 0 | $1/6$ | . | $1/6$ | $1/6$ | $d/2$ |
| G/- | 0 | 0 | 0 | 0 | 0 | 0 | 0 | 0 | 0 | 0 | $1/6$ | $1/6$ | . | $1/6$ | $d/2$ |
| T/- | 0 | 0 | 0 | 0 | 0 | 0 | 0 | 0 | 0 | 0 | $1/6$ | $1/6$ | $1/6$ | . | $d/2$ |
| - | 0 | 0 | 0 | 0 | 0 | 0 | 0 | 0 | 0 | 0 | 0 | 0 | 0 | 0 | . |

**Table S4: Summary of fractions of predicted genotypes by DelSIEVE and SIEVE for three analyzed real datasets.** Entries marked with NA denote that the corresponding method does not call the specific genotype.

| | | - | 1/- | 0/- | 0/0 | 0/1 | 1/1 | $1/1'$ |
|---|---|---|---|---|---|---|---|---|
| TNBC16 | DelSIEVE | 0 | 11.51% | 0.07% | 15.58% | 69.82% | 2.99% | 0.03% |
| | SIEVE | NA | NA | NA | 15.54% | 75.11% | 9.30% | 0.05% |
| CRC28 | DelSIEVE | 0 | 0.15% | 0.02% | 25.02% | 74.59% | 0.16% | 0.06% |
| | SIEVE | NA | NA | NA | 25.02% | 74.64% | 0.28% | 0.06% |
| CRC48 | DelSIEVE | 0 | 0 | 0.08% | 59.61% | 40.17% | 0.14% | 0 |
| | SIEVE | NA | NA | NA | 59.48% | 40.50% | 0.02% | 0 |

### References


48. Lewis, P. O. A Likelihood Approach to Estimating Phylogeny from Discrete Morphological Character Data. *Systematic Biology* **50,** 913–925. https://doi.org/10.1080/106351501753462876 (Nov. 2001).

49. Leaché, A. D., Banbury, B. L., Felsenstein, J., de Oca, A. n.-M. & Stamatakis, A. Short Tree, Long Tree, Right Tree, Wrong Tree: New Acquisition Bias Corrections for Inferring SNP Phylogenies. *Systematic Biology* **64,** 1032–1047. https://doi.org/10.1093/sysbio/syv053 (July 2015).

50. Felsenstein, J. Phylogenies from restriction sites: a maximum-likelihood approach. *Evolution* **46,** 159–173 (1992).

51. Felsenstein, J. Evolutionary trees from DNA sequences: A maximum likelihood approach. *Journal of Molecular Evolution* **17,** 368–376. https://doi.org/10.1007/BF01734359 (Nov. 1981).

52. Drummond, A. J., Nicholls, G. K., Rodrigo, A. G. & Solomon, W. Estimating Mutation Parameters, Population History and Genealogy Simultaneously From Temporally Spaced Sequence Data. *Genetics* **161,** 1307–1320. https://www.genetics.org/content/161/3/1307 (2002).

53. Bishop, C. M. & Nasrabadi, N. M. *Pattern recognition and machine learning* (Springer, 2006).

54. O'Reilly, J. E. & Donoghue, P. C. The efficacy of consensus tree methods for summarizing phylogenetic relationships from a posterior sample of trees estimated from morphological data. *Systematic Biology* **67,** 354–362 (2018).

55. Rambaut, A., Drummond, A. J., Xie, D., Baele, G. & Suchard, M. A. Posterior summarization in Bayesian phylogenetics using Tracer 1.7. *Systematic Biology* **67,** 901–904 (2018).

56. Posada, D. CellCoal: Coalescent Simulation of Single-Cell Sequencing Samples. *Molecular Biology and Evolution* **37,** 1535–1542. https://doi.org/10.1093/molbev/msaa025 (Feb. 2020).

57. Kuhner, M. K. & Felsenstein, J. A simulation comparison of phylogeny algorithms under equal and unequal evolutionary rates. *Molecular Biology and Evolution* **11,** 459–468. https://doi.org/10.1093/oxfordjournals.molbev.a040126 (May 1994).

58. Robinson, D. & Foulds, L. Comparison of phylogenetic trees. *Mathematical Biosciences* **53,** 131–147. https://www.sciencedirect.com/science/article/pii/0025556481900432 (1981).

59. Schliep, K., Potts, A. J., Morrison, D. A. & Grimm, G. W. Intertwining phylogenetic trees and networks. *Methods in Ecology and Evolution* **8,** 1212–1220. https://besjournals.onlinelibrary.wiley.com/doi/abs/10.1111/2041-210X.12760 (2017).

60. Douglas, J., Zhang, R. & Bouckaert, R. Adaptive dating and fast proposals: Revisiting the phylogenetic relaxed clock model. *PLOS Computational Biology* **17,** 1–30. https://doi.org/10.1371/journal.pcbi.1008322 (Feb. 2021).

61. Wang, K., Li, M. & Hakonarson, H. ANNOVAR: functional annotation of genetic variants from high-throughput sequencing data. *Nucleic Acids Research* **38,** e164–e164. https://doi.org/10.1093/nar/gkq603 (July 2010).

62. R Core Team. *R: A Language and Environment for Statistical Computing* R Foundation for Statistical Computing (Vienna, Austria, 2023). https://www.R-project.org/.

63. Yu, G., Smith, D. K., Zhu, H., Guan, Y. & Lam, T. T.-Y. ggtree: an r package for visualization and annotation of phylogenetic trees with their covariates and other associated data. *Methods in Ecology and Evolution* **8,** 28–36. https://besjournals.onlinelibrary.wiley.com/doi/abs/10.1111/2041-210X.12628 (2017).

64. Gu, Z., Eils, R. & Schlesner, M. Complex heatmaps reveal patterns and correlations in multidimensional genomic data. *Bioinformatics* **32,** 2847–2849. https://doi.org/10.1093/bioinformatics/btw313 (May 2016).

65. Patil, I. Visualizations with statistical details: The 'ggstatsplot' approach. *Journal of Open Source Software* **6,** 3167. https://doi.org/10.21105/joss.03167 (2021).
